# Arterial spin labelling perfusion MRI analysis for the Human Connectome Project Lifespan Ageing and Development studies

**DOI:** 10.1101/2024.09.19.613907

**Authors:** Thomas F. Kirk, Flora A. Kennedy McConnell, Jack Toner, Martin S. Craig, Davide Carone, Xiufeng Li, Yuriko Suzuki, Timothy S. Coalson, Michael P. Harms, Matthew F. Glasser, Michael A. Chappell

## Abstract

The Human Connectome Project Lifespan studies cover the Development (5-21) and Aging (36-100+) phases of life. Arterial spin labelling (ASL) was included in the imaging protocol, resulting in one of the largest datasets collected to-date of high spatial resolution multiple delay ASL covering 3,000 subjects. The HCP-ASL minimal processing pipeline was developed specifically for this dataset to pre-process the image data and produce perfusion estimates in both volumetric and surface template space. Applied to the whole dataset, the outputs of the pipeline revealed significant and expected differences in perfusion between the Development and Ageing cohorts. Visual inspection of the group average surface maps showed that cortical perfusion often followed cortical areal boundaries, suggesting differential regulation of cerebral perfusion within brain areas at rest. Group average maps of arterial transit time also showed differential transit times in core and watershed areas of the cerebral cortex, which are useful for interpreting haemodynamics of functional MRI images. The pre-processed dataset will provide a valuable resource for understanding haemodynamics across the human lifespan.

## 1. Introduction

The Human Connectome Project (HCP) Lifespan studies extend the original HCP (collected in young-adults) to characterize brain organization and connectivity during the development (HCD, 5-21 years) (Somerville et al., 2018) and ageing (HCA, 36-100+ years) (Bookheimer et al., 2019) phases of life. Taken together, the studies acquired comprehensive neuroimaging datasets for more than 3,200 individuals to facilitate characterization of the organization and connectivity of the brain at various life stages, including childhood, puberty, middle age, menopause, later-life and the ‘oldest old’. The Lifespan studies included multiple delay arterial spin labelling (ASL) MRI as an imaging modality, which permits, for the first time, perfusion measurements to be included in the HCP data releases.

ASL is a non-invasive imaging technique that uses magnetically-labelled blood as an endogenous tracer; when acquired at multiple delays (as was the case here), it permits measurement of both cerebral perfusion (CBF) and arterial transit time (ATT). Cerebral perfusion is known to vary substantially during early life (Avants et al., 2015; Satterthwaite et al., 2014), to decline with age (Kiely et al., 2022; Leidhin et al., 2021), and to be affected by many of the diseases common in older age (Alosco et al., 2013; Binnewijzend et al., 2016; Hu et al., 2010; Melzer et al., 2011; Ortapamuk & Naldoken, 2006; Patel & Markus, 2011; Prohovnik et al., 2007; Shi et al., 2016). Though ASL has been acquired in previous population studies such as the UK Biobank (Miller et al., 2016), TILDA (Donoghue et al., 2018), GENFI (Mutsaerts et al., 2019), PREVENT (Mak et al., 2021), and Whitehall II Imaging Sub-study (Suri et al., 2019); the HCP Lifespan ASL dataset is unique in its size and quality, in particular due to the use of an unusually high spatial resolution multiple delay acquisition.

When the HCP was launched in 2010, the approach adopted was novel: to acquire a large and high-quality dataset, to maintain that quality during minimal pre-processing, and then release the outputs for others to analyse further. Many aspects of this ‘HCP-Style’ to acquisition and analysis have been adopted by other studies (Glasser, Smith, et al., 2016); importantly, it has led to improvements in the reproducibility and transparency of research because all parties can access the underlying data (Elam et al., 2021). The objective of the HCP-ASL pipeline is to provide similarly high-quality pre-processed perfusion measures that others may use for downstream analysis, and many of the design decisions have been made with equivalence to existing HCP pipelines in mind in the hope of exploiting the data quality to its full extent, particularly with regards to cross-subject alignment of cortical areas.

A notable feature of existing HCP pipelines is the use of surface-based analysis to better identify functional areas of cortex and improve inter-subject registration (Coalson et al., 2018; Glasser et al., 2013; Glasser et al., 2016). In keeping with the existing HCP pipelines, the HCP-ASL pipeline produces perfusion estimates in a surface-based representation so that surface techniques may be used in downstream analysis. As few studies have yet explored surface representation of ASL-derived measures (examples include (Taso et al., 2021; Verclytte et al., 2015)), the HCP-ASL pipeline will enable the community to examine the utility of such an approach, in combination with other functional data also on the surface. The pipeline also produces volumetric outputs for compatibility with conventional analysis workflows.

This work details the processing steps performed by the HCP-ASL minimal processing pipeline and presents summary perfusion statistics calculated across the HCP Lifespan cohorts. There are numerous strategies that may be used for processing ASL data, particularly multi-delay data, and there remains debate as to which approach is best (Alsop et al., 2015; Fan et al., 2024; Lindner et al., 2023; Woods et al., 2024). We have not tried to systematically resolve these differences in this work, and instead present the pipeline as a best-effort for the unique nature of the dataset. For an example analysis of perfusion measurements derived from the data, the reader is referred to Juttukonda et al. (2021).

## 2. Materials and methods

The HCP Lifespan ASL data were processed with the HCP-ASL pipeline version 0.1.2, available at https://github.com/physimals/hcp-asl/releases/tag/v0.1.2. The following sections detail the data acquisition and the processing steps performed by the pipeline.

### 2.1. Data acquisition

ASL data for the HCP Lifespan studies was acquired in 5.5 mins of scanning on five 3T Siemens Prisma scanners at four sites (Harms et al., 2018). Data were collected for 1,306 subjects for HCD (655 female, ages 5-21 years) and 1,199 for HCA (681 female, ages 36- 102); the distribution across sites is given in Supplementary Table 1. Ethics approvals for the data collection was granted by the Washington University in St. Louis Institutional Review Board (HCA IRB ID #: 201603117, HCD IRB ID#: 201603135).

Pseudo-continuous ASL (PCASL) was used, with a 1.5s label duration and multiple post-labelling delays (PLDs) of 0.2, 0.7, 1.2, 1.7 and 2.2s, repeated 6, 6, 6, 10, and 15 times, respectively. A simultaneous multi-slice (SMS) 2D EPI acquisition with a multi-band factor of 6 was used without background suppression to achieve 2.5mm isotropic resolution (2.27mm slice thickness plus 10% gap), with 60 slices total, 59ms readout time per slice and 19ms echo time (further details are given in (Li et al., 2015)). Although 2D readouts have inherently lower signal-to-noise ratio than 3D, they do not introduce substantial spatial blurring due to T2 decay during the readout (Alsop et al., 2015; Woods et al., 2024). Such blurring would reduce the accuracy of registration between the ASL data and T1w anatomical image, negatively affecting the surface parameter maps that are the desired endpoint of the pipeline. Blurring can be reduced via the use of segmented 3D acquisitions, but such acquisitions are highly sensitive to motion. For this reason, after comparing 3D and 2D (with SMS) PCASL, HCP Lifespan adopted the latter (Harms et al., 2018).

To calibrate perfusion measurements into units of ml/100g/min, two 2.5mm isotropic PD-weighted M0 calibration images (TR>8s) were acquired at the end of the PCASL scan. In addition, a strong pre-saturation covering the imaging region was played out just before the PCASL module using 3 selective RF pulses with the spoiling gradients for each applied along two different directions (i.e., the first pre-saturation pulse was followed with spoiling gradients along the readout and slice directions; the second along the phase and slice directions; and the third along the readout and phase directions). For susceptibility distortion correction, two phase-encoding-reversed spin-echo images were acquired that were geometrically and distortion matched to the ASL data. T1 and T2-weighted anatomical and functional MRI images were acquired and pre-processed via the existing HCP minimal processing pipelines (Glasser et al., 2013, 2018, 2019; Harms et al., 2018; Robinson et al., 2018).

### 2.2. Data pre-processing and corrections

The ASL images contained several geometric and intensity artefacts that required correction before perfusion estimation could be performed. The use of a pre-saturation pulse in a multi-slice acquisition scheme contributed to slice-wise intensity variations in the inferior-superior direction in the ASL timeseries and calibration images, most notably between the last and first slices of adjacent bands, illustrated in Figure 1.

**Figure 1:**
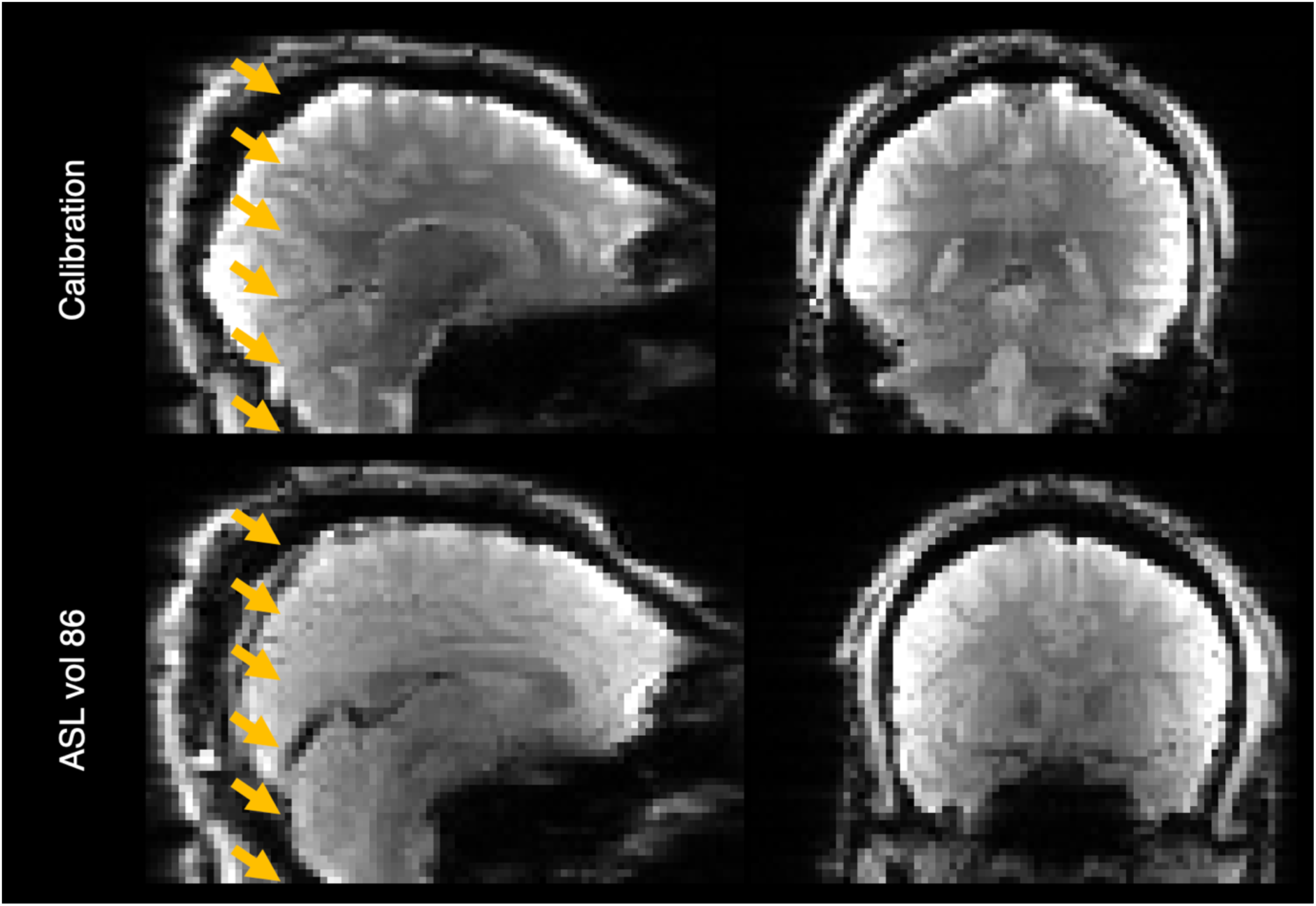
Banding artefact visible on the calibration image (top row) and last volume of the ASL timeseries (bottom row). The bottom of each band (six in total) is shown with an arrowhead.

The perfusion signal in ASL is obtained by subtracting successive pairs of label and control images in the ASL timeseries. This means that ASL is sensitive to within-timeseries motion which can lead to spurious signal when mis-aligned images are subtracted, and it is therefore necessary to motion-correct the data. Conventional methods for rigid-body motion correction such as FSL MCFLIRT (Jenkinson et al., 2002) may not be accurate when applied to banded data because the registration cost function may reward alignment of the intensity banding rather than anatomy, leading to anatomical misalignment, which will produce spurious signal after label-control subtraction. Furthermore, even if there is perfect anatomical alignment within a label-control pair, in the presence of motion there will be artefactual intensity variations due to the bands occurring at different anatomical locations across the pair, resulting in imperfect elimination of the banding effects of static tissue signal. It was therefore decided to “de-band” the data before performing motion correction.

The first banding correction applied to the data accounted for the different relaxation times of each slice in the 2D readout, which gives rise to a saturation recovery effect in the signal. While this effect exists in a conventional non-SMS 2D ASL acquisition (Golay et al., 2004), with SMS acquisitions there are adjacent slices within brain tissue acquired in the first vs. last SMS excitation, making the discontinuity more evident, and importantly, contributing to the prominent banding artifact that was observed. To remove the saturation recovery effect, a simplified model of the form 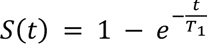 was fitted to the control images of the ASL timeseries using FSL FABBER, where *T1* is the longitudinal relaxation time of tissue and *t* the slice-specific interval from the pre-saturation module (which is the sum of the duration of the labelling module, the PLD, and the slice-specific delay in the readout module, accounting properly for the SMS nature of the acquisition). An example T1 map generated by FABBER is given in Supplementary Figure 1. The saturation recovery differences were then corrected (in both the control and label images) by normalizing the intensities to the estimated value that they would have had if all the slices had been acquired simultaneously at the PLD.

However, during pipeline development, it was found that saturation recovery correction alone was unable to fully remove banding in both the calibration image and later PLDs of the ASL timeseries (illustrated in the results section). Given the long TR of the calibration image (8s), or long recovery times of the later PLDs (up to 3.7s), theory predicts that longitudinal magnetisation will have largely recovered following those intervals and therefore the saturation recovery effect will be minimal. The presence of banding in the calibration images and later PLDs implied that there was a second banding mechanism (possibly a magnetisation transfer effect, addressed in the Discussion), for which a correction was empirically derived using the calibration images of 80 subjects drawn equally from the HCA and HCD cohorts. After performing saturation recovery correction and masking the images to include grey and white matter only, a linear model for slice-wise mean intensity was fitted to each band and the coefficients averaged across subjects (see Supplementary Material for details).

Correction for gradient nonlinearity distortion was performed using *gradunwarp* operating on the calibration image with the gradient coefficients file for the Prisma system (as used in the HCP *PreFreeSurfer* pipeline (Glasser et al., 2013)). Susceptibility distortion correction was performed using FSL TOPUP operating on a pair of phase-encode reversed spin-echo images that had undergone gradient distortion correction. Bias field (B1) estimation was performed using the spin echo-based approach (‘SEBASED’, Glasser et al., 2016) operating on the calibration image that had undergone gradient distortion, susceptibility distortion and banding correction.

Figure 2 shows how the aforementioned preprocessing corrections were derived and combined in the HCP-ASL pipeline to produce the fully corrected ASL and the calibration images on which perfusion estimation can be performed. Though two calibration images were acquired, only the first was used in the pipeline, though both were fully corrected. Some corrections were refined iteratively: for example, given that banding is expected to have a strong influence on the accuracy of motion estimation, each was performed twice on the assumption that an improvement in one should lead to an improvement in the other. Motion estimation was performed using MCFLIRT with the calibration image as the reference to yield a registration between the calibration image and ASL timeseries. In total, four subject specific inputs and two generic inputs to the pipeline are required: for the individual, the phase encode reversed fieldmaps, the calibration image, the T1w structural image and the ASL timeseries; and generically, the Siemens Prisma gradient coefficients and the empirical banding correction coefficients.

**Figure 2:**
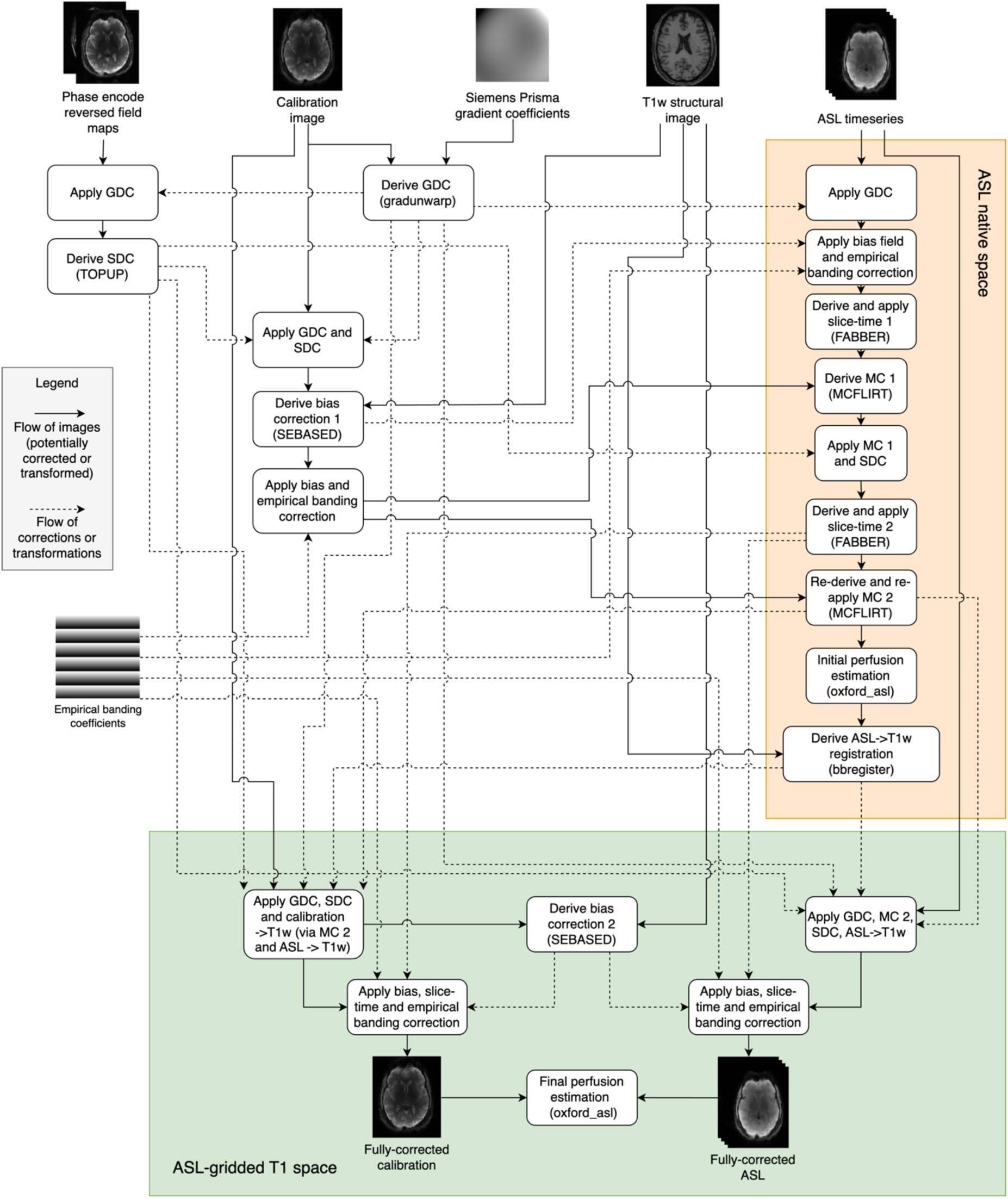
Schematic diagram showing how the fully corrected calibration and ASL images in ASL-gridded T1 space are derived (green box). Note that some intermediate registrations and transformations are not shown for clarity (particularly in the orange shaded box); all operations applied to the fully corrected outputs are shown. SEBASED: spin-echo based, SDC: susceptibility distortion correction, GDC: gradient distortion correction, MC: motion correction, FABBER: FSL model-fitting tool, MCFLIRT: FSL motion correction tool.

**Figure 3:**
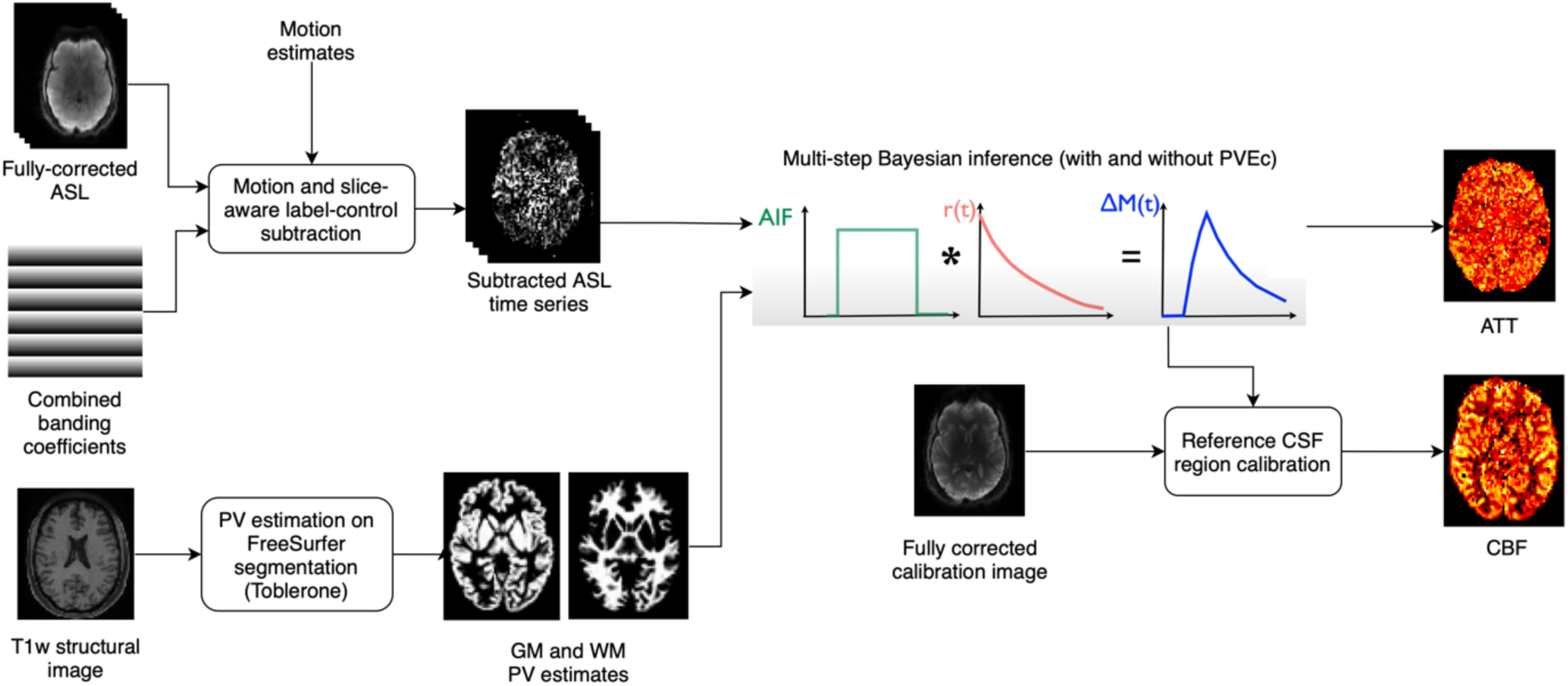
Perfusion analysis applied to ASL difference data after alignment with the individual’s T1w image. Bayesian inference is used to produce both partial volume corrected and non-corrected estimates of perfusion and arterial arrival times, as well as arterial cerebral blood volume estimates. During inference the slice timing image on the right is used to produce slice specific PLDs. Calibration of perfusion measures was performed using CSF reference region calibration.

### 2.3. Data alignment within individuals

In keeping with HCP fMRI pipelines, the HCP-ASL pipeline transforms ASL timeseries data into alignment with the structural images before analysis (Glasser et al., 2013), though retains the original 2.5mm resolution (hereafter referred to as “ASL-gridded T1 space”). Multiple steps were used to obtain an accurate ASL to T1w registration. An initial registration was made using FreeSurfer’s boundary-based registration (BBR, bbregister) between the calibration image and the T1w image (Greve & Fischl, 2009; Jenkinson et al., 2002); this was combined with the motion correction matrices to obtain a registration between ASL and T1w. The calibration image has better grey-white tissue contrast than the un-subtracted ASL timeseries which facilitates accurate BBR. Then, perfusion estimation on the native ASL data with all banding and distortion corrections was performed to obtain a CBF map with increased grey-white tissue contrast, on which a second BBR was performed to the T1w image. The CBF map from this step was then discarded.

The transform between the ASL and calibration images and T1w alignment was merged with the motion correction, susceptibility distortion and gradient distortion transforms and applied in a single step to the raw ASL and calibration images using the *regtricks* library to produce the fully-transformed ASL and calibration images in ASL-gridded T1 space (Kirk, 2022). Jacobian intensity correction for distortion-induced signal pile-up was performed and cubic splines with prefiltering used to perform the interpolation (Unser et al., 1993). Minimising the number of interpolation operations aimed to reduce the introduction of additional partial volume effects that cannot be entirely removed with subsequent correction. SEABASED bias field estimation was re-run on the fully-transformed calibration image to obtain bias field correction coefficients in ASL-gridded T1 space.

To intensity correct (banding and bias) the fully-transformed ASL and calibration images, the calibration to T1w registration was applied to the empirical banding coefficients, and the ASL to T1w registration to the saturation recovery correction coefficients, to transform both banding corrections into ASL-gridded T1 space. Given the ASL and calibration image data were distortion corrected, no distortion correction was applied to the banding correction coefficients (it is only necessary to correct one or the other). The empirical banding coefficients were combined with the newly-derived bias field estimates to intensity correct the calibration image, and both sets of coefficients were combined with the bias field estimates to intensity correct the ASL image.

### 2.4. Perfusion analysis

In the presence of head motion, voxels travelling between neighbouring slices during the acquisition will receive differing intensity scaling during the banding corrections, which could lead to spurious signal after label-control subtraction. The general linear model (GLM) framework for motion-aware subtraction of banded and background-suppressed ASL data developed by Suzuki et al. (2019) was used in this work, with the small modification that it was not necessary to account for background suppression because it was not used during the image acquisition. The subtracted timeseries data was then used for perfusion estimation.

CBF and ATT estimation were performed using a variational Bayesian method via the *oxford_asl* script (Chappell et al., 2009, 2023). The *aslrest* Buxton model with CBF, ATT and macrovascular components was used (Buxton et al., 1998; Chappell et al., 2011). A normal distribution prior with mean 1.3s was used on ATT and an automatic relevancy determination (ARD) prior was used on macrovascular perfusion to remove this component from non-arterial voxels. Slice-timing correction was performed by adjusting the PLDs in each voxel by their slice timing offset, and perfusion was converted from arbitrary units into ml/100g/min using the mean signal value of CSF in the lateral ventricles from the calibration image (Pinto et al., 2020). Reference region calibration was used instead of a voxel-wise strategy due to the availability of a high-resolution calibration image, which allowed for accurate ventricular segmentation with minimal PVE, as well as mitigating the possibility of introducing (potentially uncorrected) banding artefacts from the calibration image if voxel-wise division were used. The tissue T1 values previously estimated for saturation recovery correction were passed to *oxford_asl*.

ASL is typically acquired with ∼4mm voxel sizes which gives rise to partial volume effects (PVE) caused by the coarse spatial resolution of the data in relation to the cerebral cortex, which has a mean thickness of around 2.5mm. There is ongoing debate as to whether PVE correction (PVEc) should be routinely used in ASL studies. Though it is theoretically justified, particularly for studies covering a wide range of participant ages, there remain questions around the robustness and efficacy of currently available methods (Chappell et al., 2021). The ASL data acquired by the Lifespan HCP had a relatively high spatial resolution (2.5mm isotropic), which not only reduced the PVE but also enabled more accurate PVEc to be performed. PVEc is performed in the voxel space using a spatial variational Bayes method implemented in *oxford_asl* (Chappell et al., 2011); the required partial volume estimates are obtained using Toblerone operating on the FreeSurfer-derived cortical surfaces and subcortical segmentations (Fischl, 2012; Kirk et al., 2020). For PVEc, a normal distribution prior with mean of 1.3s was used for GM ATT and 1.6s for WM ATT.

Both PVEc and non-PVEc perfusion estimates are generated so that the end-user may decide which is most appropriate for their application. For non-PVEc, spatial regularisation is not used (deviating from the recommended settings for *oxford_asl*, which were otherwise used) to minimise the mixing of signal across the grey/white cortical boundary, at the cost of somewhat greater image noise. By contrast, for PVEc, spatial regularisation is an intrinsic feature of the spatial variational Bayes method used by *oxford_asl*.

### 2.5. Output in standard space; calculation of Image-Derived Phenotypes

Volumetric CBF and ATT maps from *oxford_asl* are produced in both ASL-gridded T1w space and MNI152 2mm template space, via the existing FNIRT registration produced by the HCP PreFreeSurfer structural processing pipeline. To produce output parameter maps in greyordinates space (a combined surface and volumetric space represented in CIFTI format), the volumetric outputs of *oxford_asl* are projected onto the individual’s native cortical surface using the HCP ribbon-constrained method, registered with MSMAll multi-modal areal-feature-based registration and resampled to a common surface mesh, and finally smoothed with a 2mm full-width half-maximum kernel using a surface-constrained method (Glasser et al., 2016; Glasser et al., 2013; Robinson et al., 2014, 2018). The surface and MNI outputs (masked to consider subcortical structures only) were combined to produce the final CIFTI greyordinates files. Image-derived phenotypes (IDPs) were produced for each participant by computing the mean and standard deviation within each parcel of the HCP multi-modal parcellation (version 1.0) for both PVEc and non-PVEc variants of CBF and ATT.

### 2.6. Quality control

Though the HCP-ASL pipeline does not perform any automated QC, it does produce intermediate outputs that enable researchers to perform visual checks. These are produced as a single Workbench scene file (a pre-specified visualisation) containing seven scenes, each of which is saved as a standalone image to facilitate quick inspection (although full exploration requires opening the scene in Workbench). The scenes illustrate a) the bounding box of the ASL timeseries after motion correction within the acquisition field of view (voxels missing data due to their movement outside of the acquisition field-of-view for a single timepoint are excluded from perfusion estimation) and the quality of registration to the T1w scan, illustrated in Figure 4; b) the raw ASL timeseries before and after all corrections are applied; c) non- PVEc CBF displayed on the inflated cortical surface and in subcortical structures; d) non-PVEc ATT displayed in the same manner; e) PV estimates for GM and WM; and the ventricular CSF mask used for calibration; f) a volumetric montage of non-PVEc CBF; and g) a volumetric montage of non-PVEc ATT.

**Figure 4:**
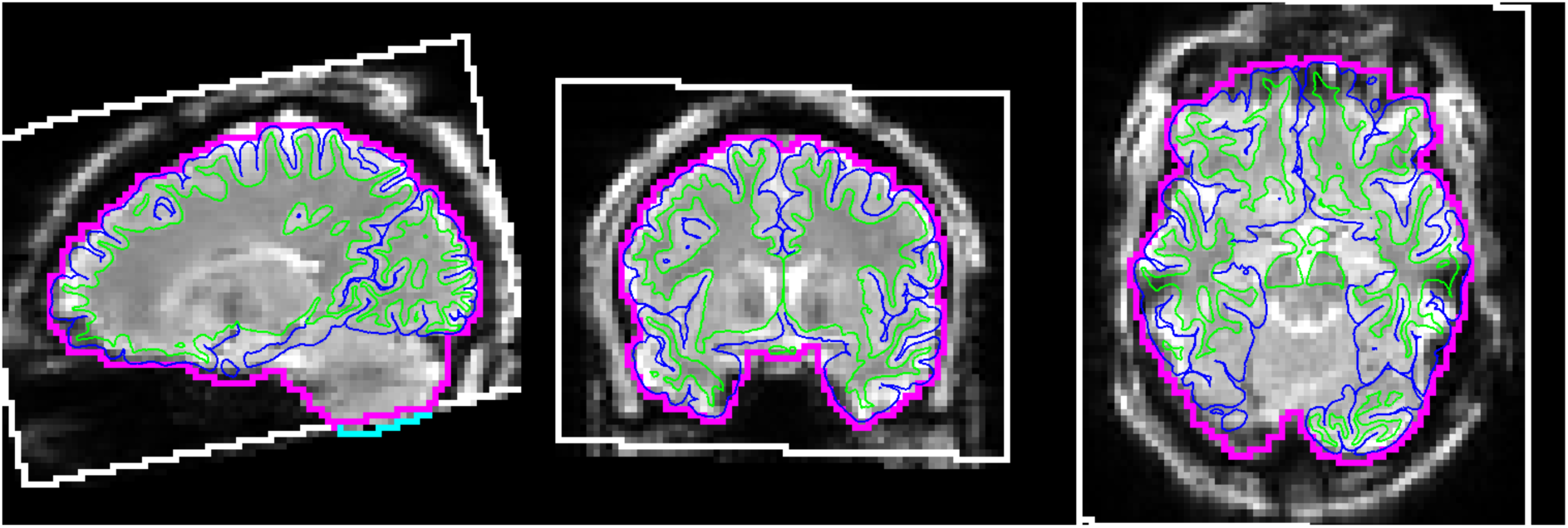
Workbench scene produced by the pipeline to assess registration and masking accuracy. The FreeSurfer white and pial surfaces are shown via thin green and blue lines, respectively. The ASL volumetric brain mask outline is shown in magenta. The white box denotes the field of view of the ASL acquisition, transformed into the ASL-gridded T1w space. The cyan line (seen at bottom of cerebellum in the sagittal view) denotes a section of the ASL brain mask that lies outside the field of view. The base image in greyscale is the first volume of the fully-corrected ASL timeseries image.

## 3. Results

### 3.1. Data pre-processing

Figure 5 shows the effect of the two individual banding corrections (empirically-derived and saturation recovery) individually and combined together on a slice-wise basis. The two banding mechanisms worked in opposing directions: at early PLDs, the saturation recovery effect was almost equal and opposite to the empirical banding effect, leading to a subtle overall correction, whereas for late PLDs where the saturation recovery effect was weaker, the empirical banding effect dominated.

**Figure 5:**
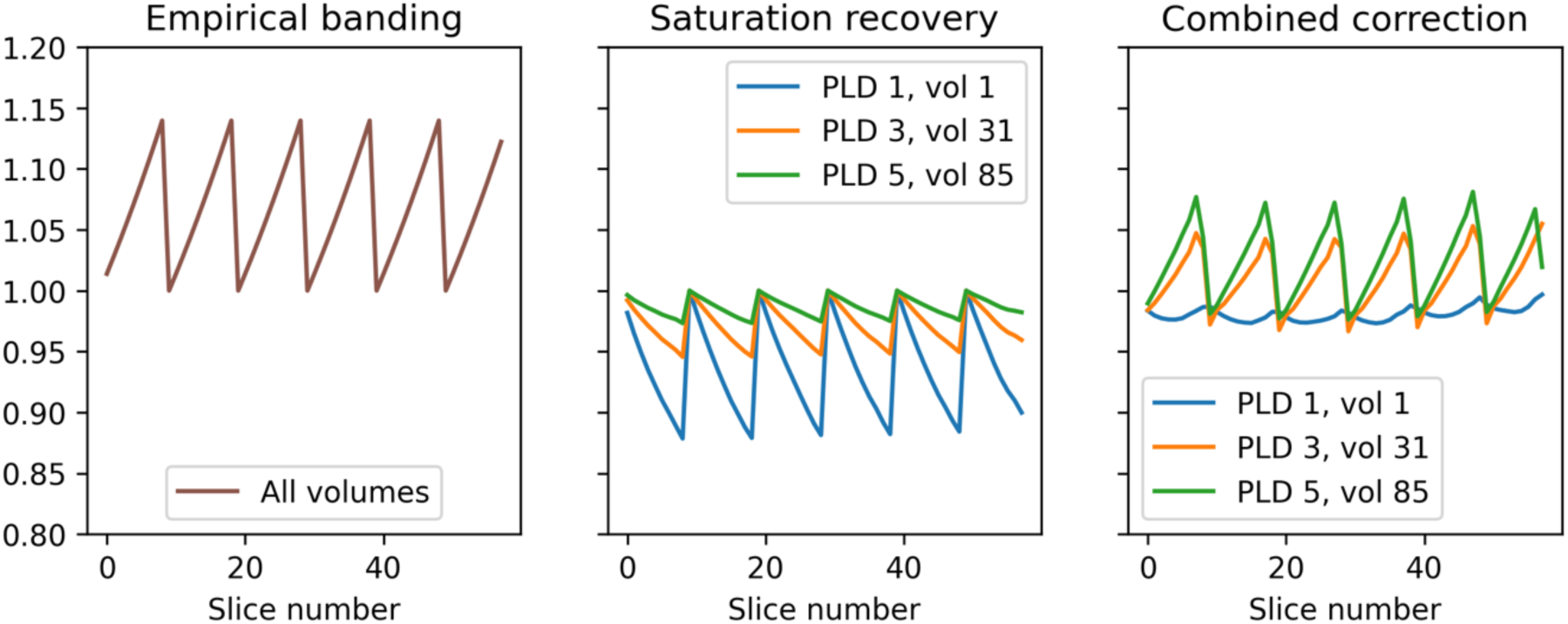
Magnitude of the two multiplicative banding corrections according to slice number and PLD/volume number within the ASL timeseries. First panel: the empirical banding correction applies equally to all PLDs/volumes within the ASL timeseries. Second panel: the saturation recovery correction is stronger for early PLDs than later ones (0.2s vs 1.2s vs 2.2s respectively) and operates in the opposite direction to the empirical banding correction. Final panel: when the two corrections are combined, they almost cancel each other in early PLDs and have a stronger effect in later PLDs (due to the reduced magnitude of the saturation recovery correction).

Figure 6 shows the effect of the two banding corrections applied in isolation and together to the ASL timeseries from participant HCD0378150. The saturation recovery correction in isolation had minimal effect on the volumes of the ASL series with the longest (2.2s) PLDs. This was consistent with theoretical prediction: at longer PLDs, the relative differences in slice-timing decrease, which will reduce the prominence of the saturation recovery effect, as was observed. Conversely, the empirical correction in isolation was unable to fully remove banding on the shortest (0.2s) PLDs, for which the saturation recovery effect is substantial. For all PLDs, the combination of the two separate corrections was better able to remove banding than in isolation.

**Figure 6:**
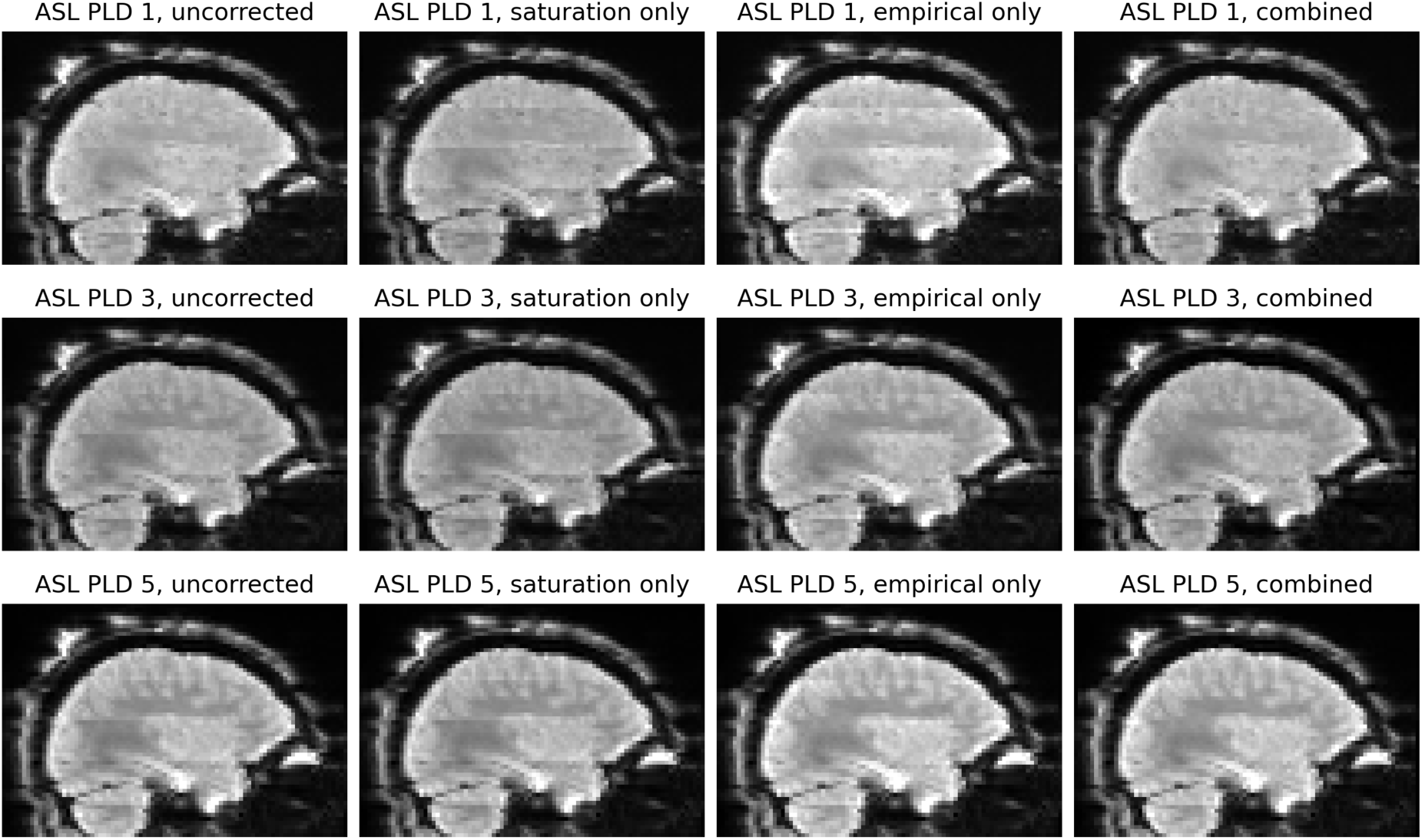
Banding artefacts visible in the first (0.2s), third (1.2s) and final (2.2s) PLDs of the ASL timeseries. First column: before correction, visible banding varies according to PLD, because the saturation recovery mechanism affects each PLD differently. Second column: saturation recovery correction in isolation is unable to remove all banding at any PLD. Third column: the empirical banding correction in isolation does not remove banding in the early PLDs but does so in later PLDs. Fourth column: at all PLDs, the combination of the two banding corrections better removes banding than in isolation. The images in all columns are single volumes from the corresponding PLD group, and were previously corrected for gradient nonlinearity distortion, susceptibility distortion, and the receive-coil bias field.

### 3.2. Group average CBF and ATT maps

Figure 7 shows the group average CBF maps for the HCA cohort, both with and without PVEc. In these and the following figures, the HCP multi-modal cortical parcellation (version 1.0) has been overlaid in black on the surface (Glasser, Coalson, et al., 2016). Areas of differential CBF (colour transitions) were observed to follow areal boundaries on the cortical surface. PVEc notably increased CBF, particularly in cortical tissue.

**Figure 7:**
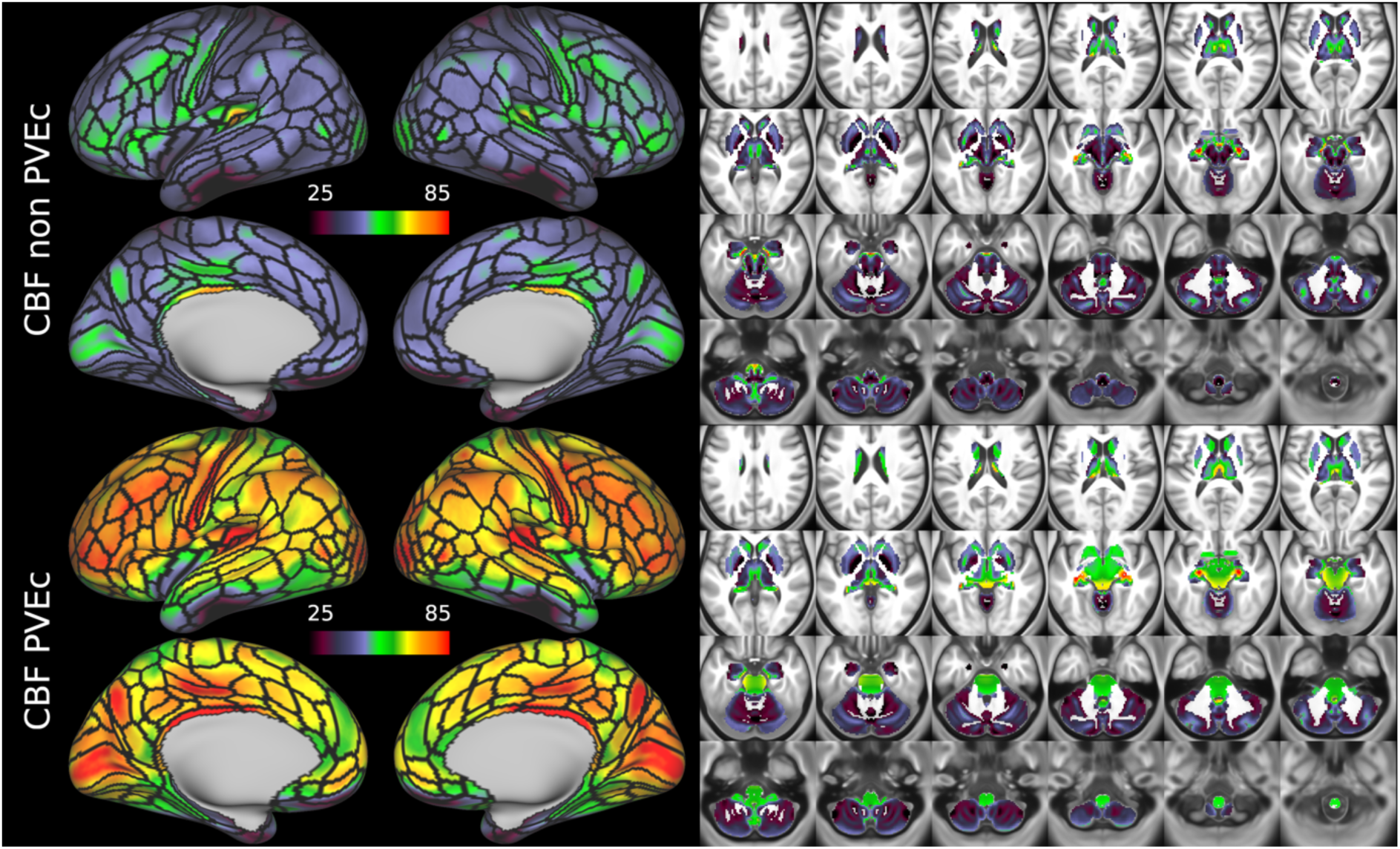
Group average CBF maps (in ml/100g/min) for the HCA cohort in the cortex and subcortical structures. PVEc led to substantial increases in CBF in the cortical ribbon; the increase was less pronounced in the subcortex.

Figure 8 shows the group average ATT maps for the HCA cohort, both with and without PVEc. Along with the volumetric representation given in Figure 11, ATT showed correspondence with known vascular anatomy, namely that the cores of the vascular territories (middle, anterior, and posterior cerebral arteries; MCA, ACA, PCA) were perfused before watershed regions that lie along the margins of these vascular territories; and anterior circulation (supplied by the MCA and ACA) was perfused before posterior circulation (supplied by the PCA). PVEc had a negligible effect on ATT.

**Figure 8:**
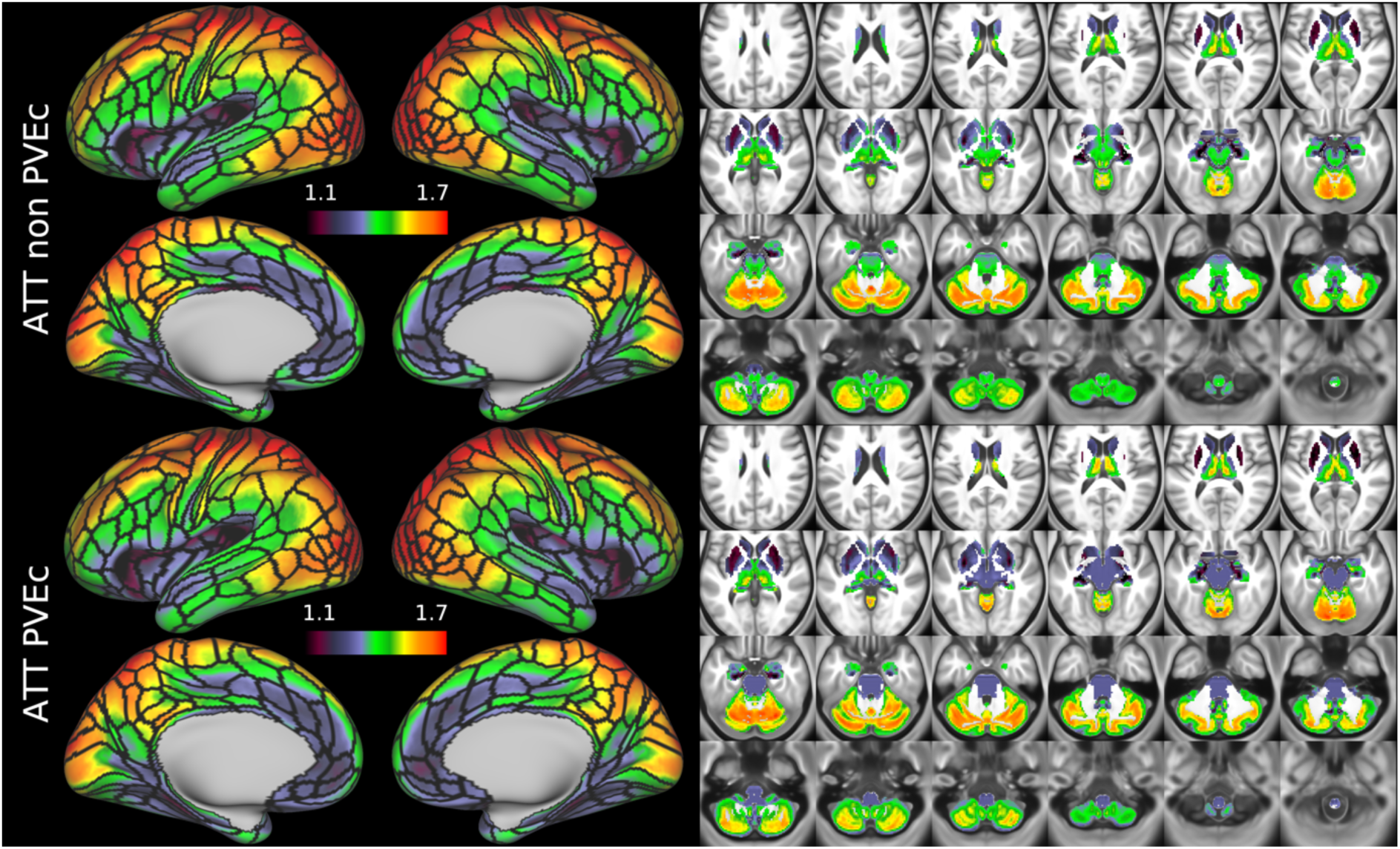
Group average ATT maps (in sec) for the HCA cohort in the cortex and subcortical structures. PVEc did not lead to notable changes in ATT.

Figure 9 and Figure 10 show group average CBF and ATT maps for the HCD cohort, both with and without PVEc. The ATT maps showed longer transit times for posterior and watershed regions.

**Figure 9:**
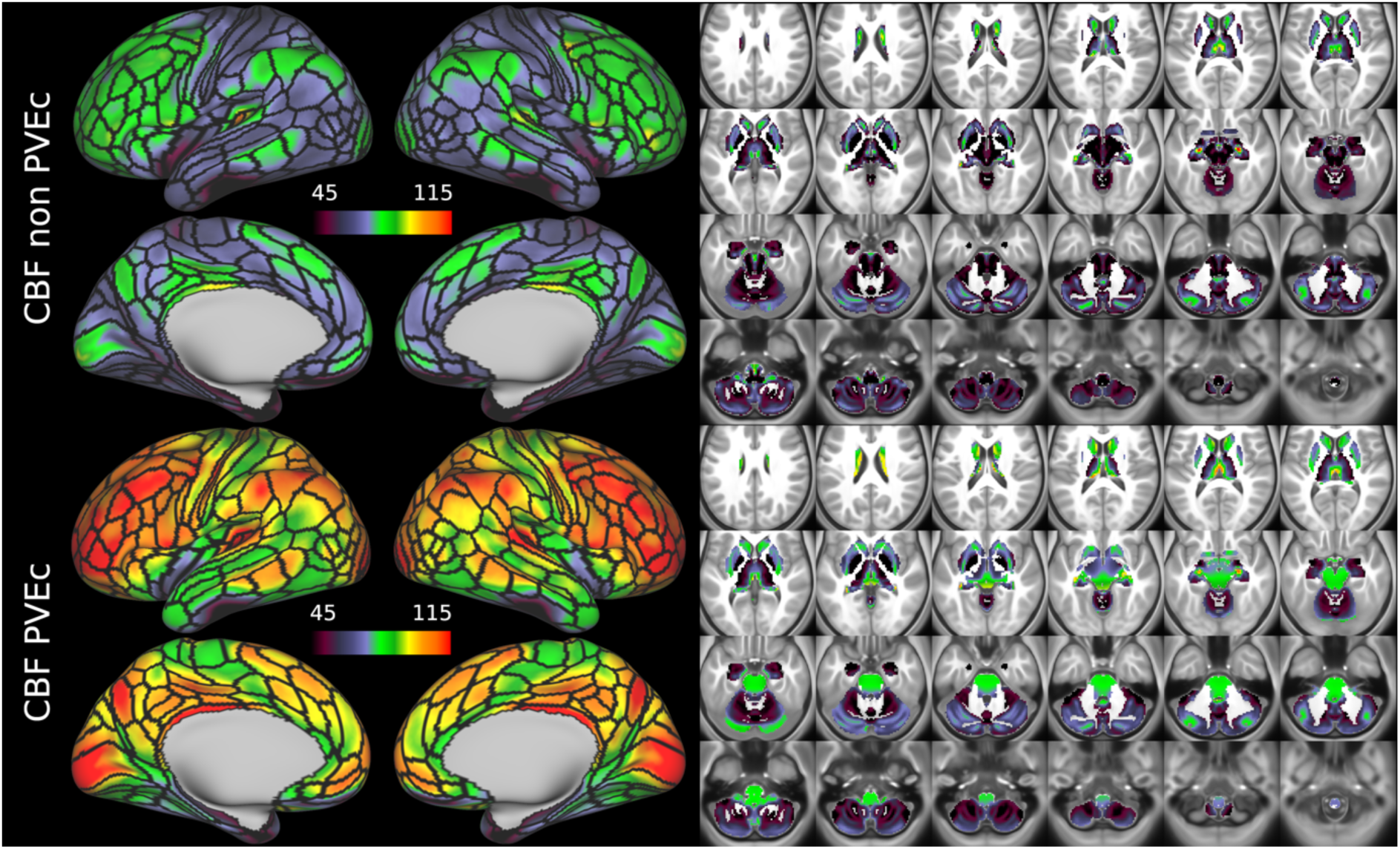
Group average CBF maps for the HCD cohort in the cortex and subcortical structures. PVEc led to substantial increases in CBF in the cortical ribbon; the increase was less pronounced in the subcortex.

**Figure 10:**
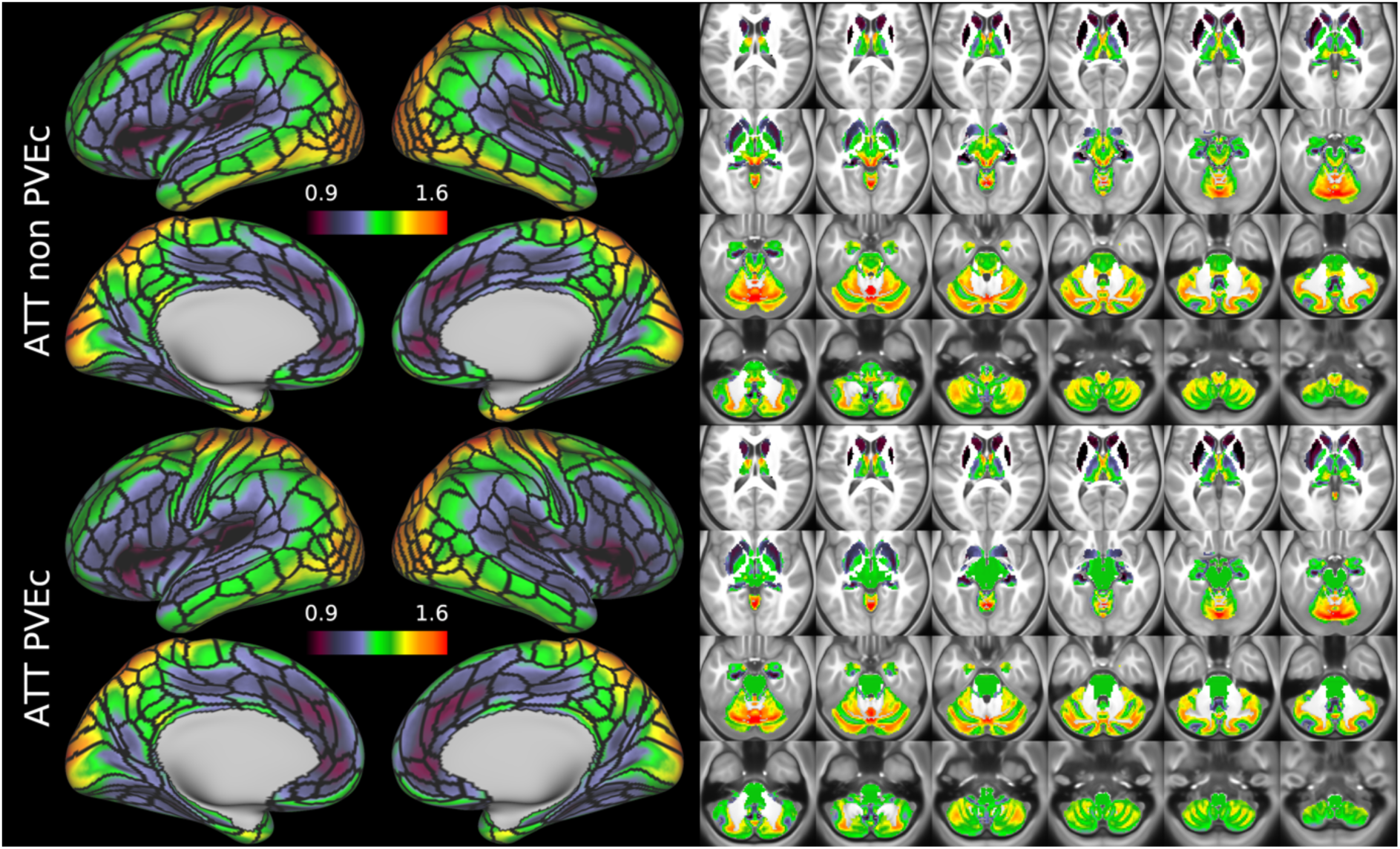
Group average ATT maps for the HCD cohort in the cortex and subcortical structures. PVEc did not lead to notable changes in ATT.

**Figure 11:**
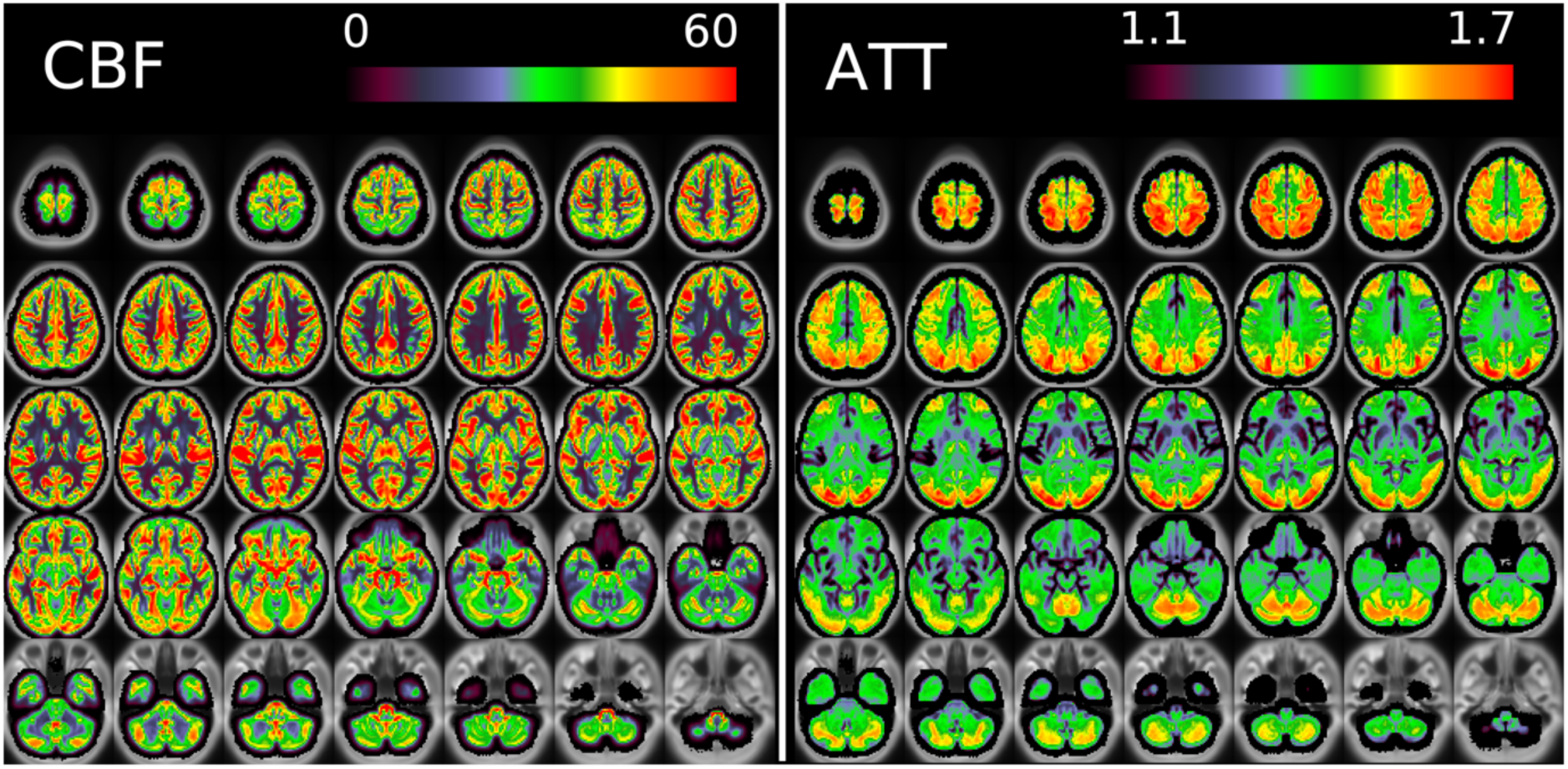
Group average CBF and ATT maps without PVEc for the HCA cohort in volumetric representation.

Figure 11 and Figure 12 show group average maps for non-PVEc CBF and ATT in each cohort, shown in a volumetric representation. This is a conventional representation of the same data shown the preceding figures.

**Figure 12:**
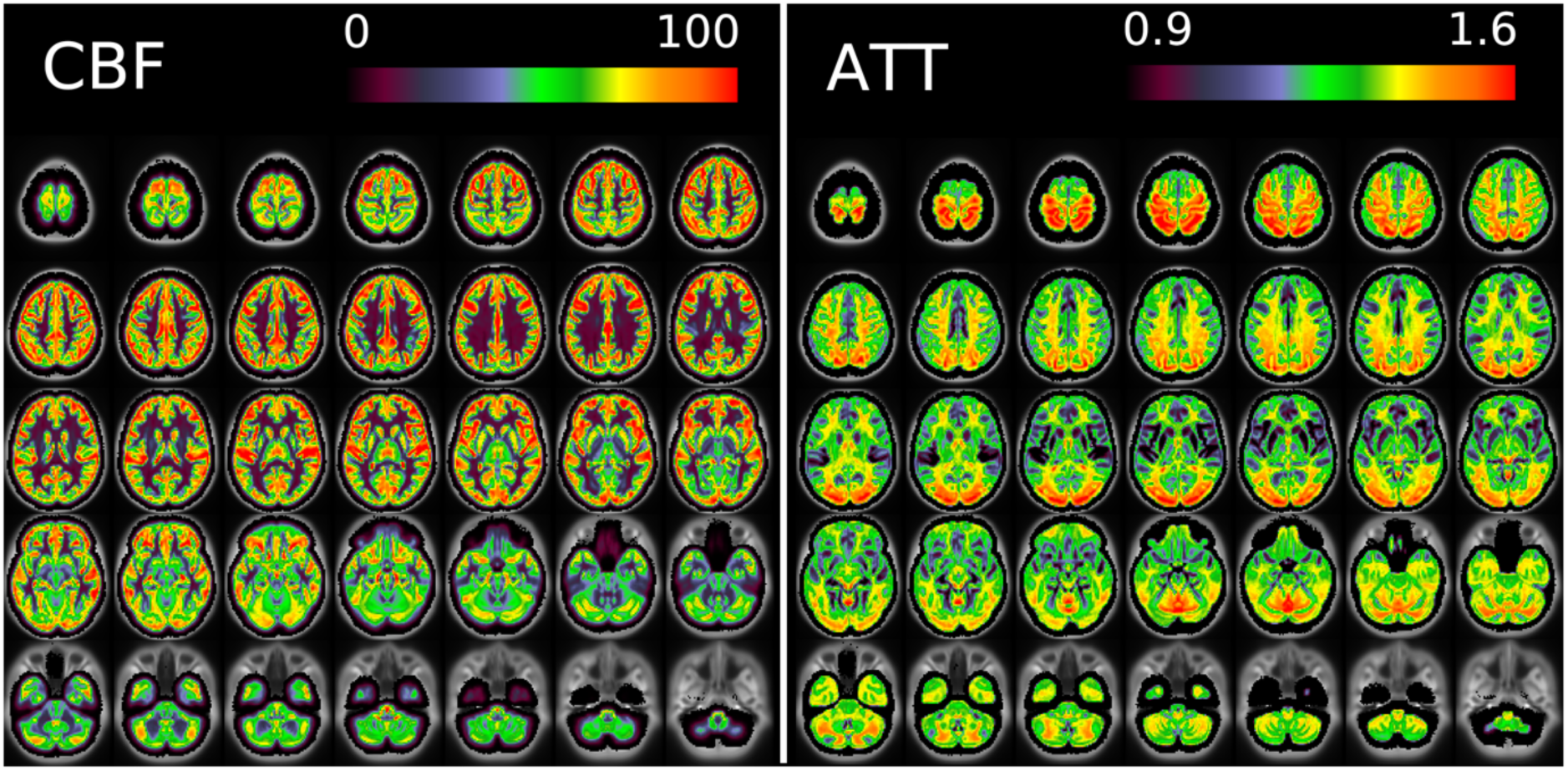
Group average CBF and ATT maps without PVEc for the HCD cohort in volumetric representation.

### 3.3. The effect of partial volume correction

In both cohorts, PVEc notably increased CBF, particularly in cortical tissue, but had a negligible effect on ATT.

Figure 13 shows the effect of PVEc on CBF and ATT measurements, separated into subcortical and cortical greyordinates, for a single subject. For cortical greyordinates, the average increase in CBF following PVEc was around 15 ml/100g/min, whereas for subcortical greyordinates it was around 5ml/100g/min. For ATT, both cortical and subcortical greyordinates showed negligible average increases.

**Figure 13:**
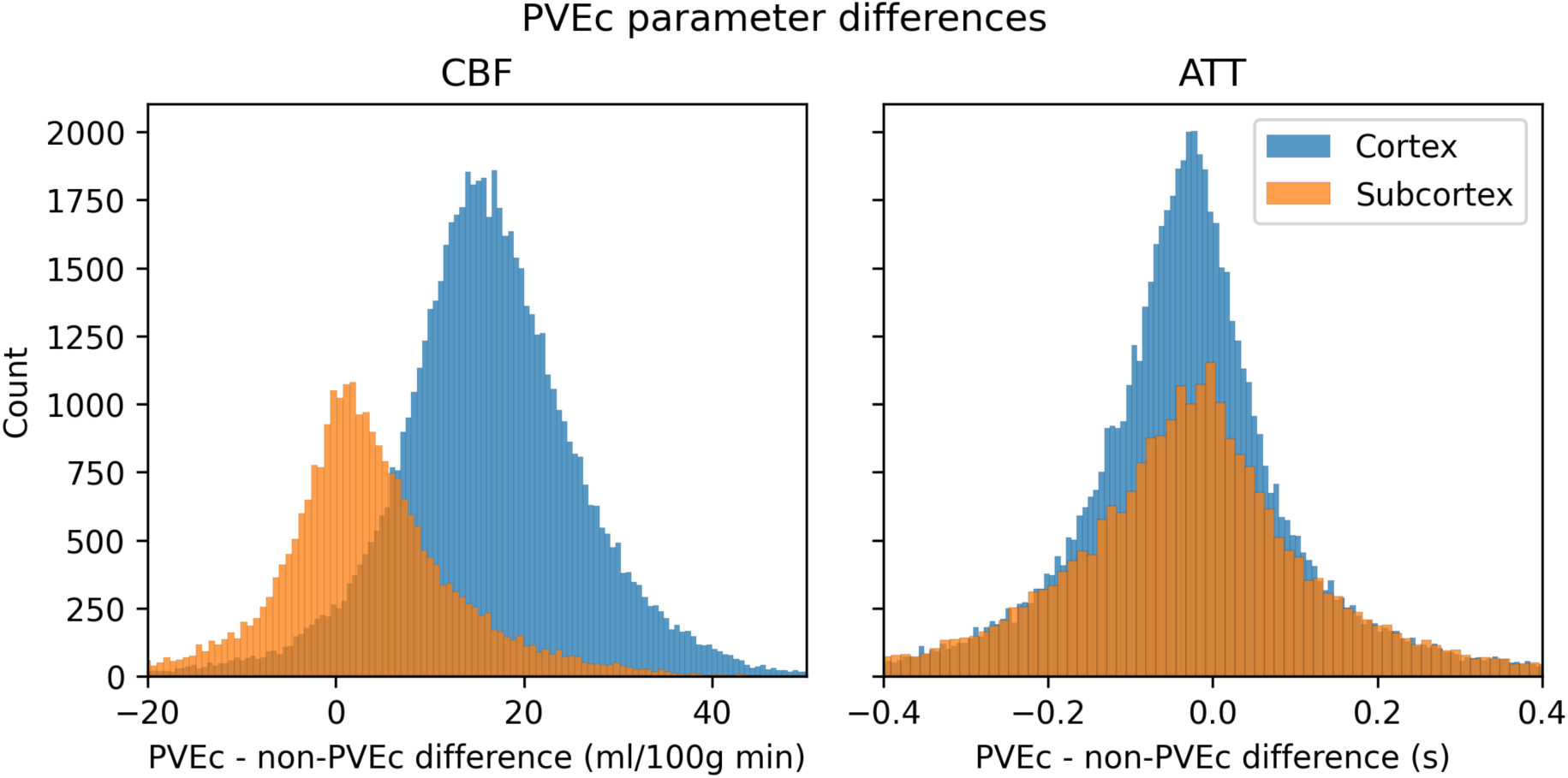
The effect of PVEc on CBF (left) and ATT (right), shown as a histogram of differences in cortical and subcortical greyordinates for a single HCD subject. In the cortex, the mean increase in CBF was around 15 ml/100g/min, whereas in the subcortex the increase was around 5 ml/100g/min. For ATT, the mean difference was close to 0s and there was no substantial difference in the distribution between cortical and subcortical greyordinates.

Figure 14 shows distributions of each subject’s mean CBF and ATT across cortical and subcortical grayordinates before and after PVEc, grouped by cohort. Numerical values for each distribution mean are given in Table 1. Between cohorts, statistically significant differences were observed (all comparisons with *p* < 0.05). On the non-PVEc data, the HCA cohort had lower CBF than HCD (45 vs 68 ml/100g/min) and longer ATT (1.46 vs 1.28s).

**Figure 14:**
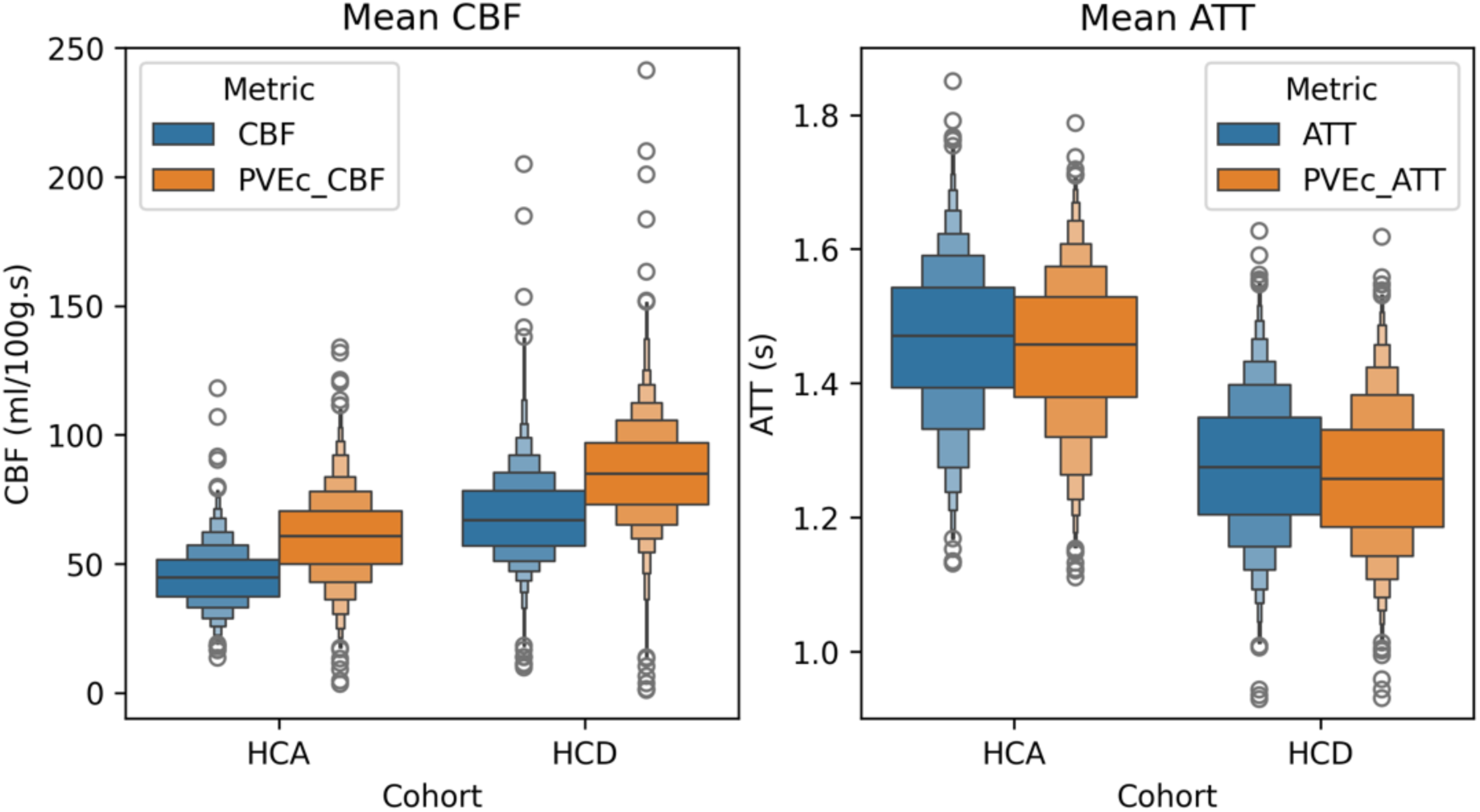
Mean GM CBF and ATT across individuals without and with PVEc for the two cohorts.

**Table 1:** Mean parameter values within cohorts. The differences between cohorts for both PVEc and non-PVEc values were statistically significant (independent t-test not assuming equal variance) in all cases. Cohen’s d was calculated for independent samples using pooled variance.

| Parameter | HCA | HCD | $\Delta$ (HCD - HCA) | t-statistic | p-value | Cohen's d |
| --- | --- | --- | --- | --- | --- | --- |
| PVEc CBF | 60.54 | 85.22 | 24.69 | 39.26 | $< 1e-10$ | 1.38 |
| CBF | 45.02 | 68.41 | 23.39 | 43.53 | $< 1e-10$ | 1.54 |
| PVEc ATT | 1.45 | 1.26 | -0.19 | -51.07 | $< 1e-10$ | -1.79 |
| ATT | 1.46 | 1.28 | -0.19 | -49.80 | $< 1e-10$ | -1.74 |

Table 2 shows that PVEc significantly increased CBF in both cohorts: from 45 to 61 ml/100g/min in HCA from 68 to 85 ml/100g/min in HCD. For both cohorts, PVEc led to a small but statistically significant decrease in ATT on the order of 0.01s.

**Table 2:** Within cohorts, PVEc lead to large changes in CBF and small changes in ATT. In all cases, the difference between PVEc and non-PVEc was statistically significant (paired t-test). Cohen’s d was calculated for paired samples as the mean of differences divided by the standard deviation of differences.

| Cohort | Parameter | Non-PVEc | PVEc | $\Delta$ (PVEc - nonPVEc) | t-statistic | p-value | Cohen's d |
| --- | --- | --- | --- | --- | --- | --- | --- |
| HCD | CBF | 68.41 | 85.22 | 16.82 | 100.98 | < 1e-10 | 2.54 |
| HCD | ATT | 1.28 | 1.26 | -0.02 | -27.72 | < 1e-10 | -0.70 |
| HCA | CBF | 45.02 | 60.54 | 15.52 | 102.44 | < 1e-10 | 2.51 |
| HCA | ATT | 1.46 | 1.45 | -0.01 | -20.15 | < 1e-10 | -0.49 |

### 3.4. Individual subject CBF and ATT maps

Figure 15 shows CBF and ATT maps for a single participant of the HCD cohort. Although the individual subject maps were noisier than the group average, it was possible to observe similarities with the group average maps; for example, elongated ATT in posterior and superior regions of cortex.

**Figure 15:**
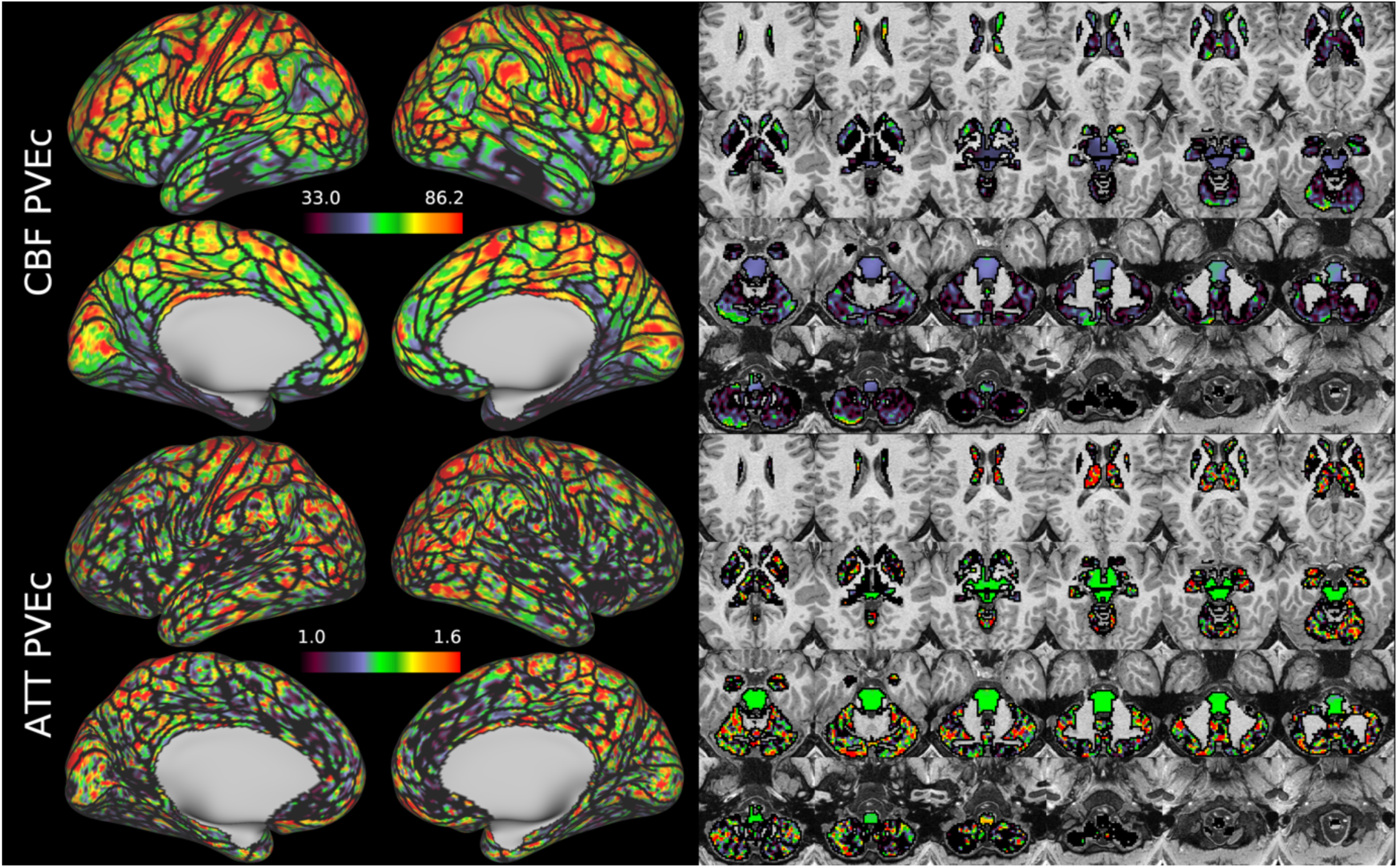
CBF and ATT maps (with PVEc) in the cortex and subcortical structures for subject HCD0378150 of the HCD cohort.

Figure 16 shows the same HCD individual as Figure 15, where CBF and ATT have been averaged within the parcels of the MCP multi-modal parcellation. This reduced the dimensionality of the data dramatically, enabling the data in Figure 12 to be represented in 379 parcels (whereas there are 91282 greyordinates in Figure 11).

**Figure 16.**
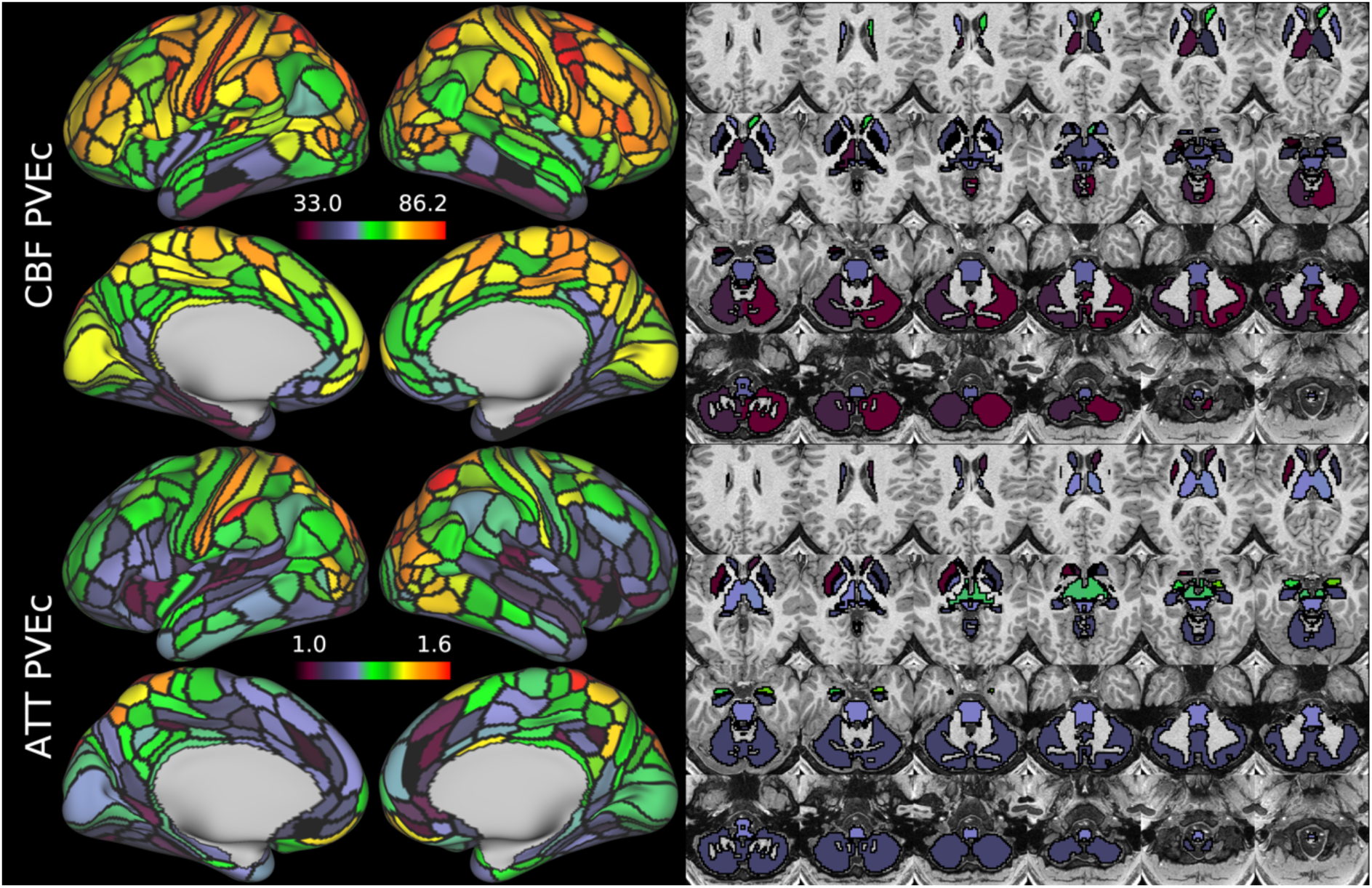
shows the same individual and measures as Figure 15 represented as IDPs using the HCP multi-modal parcellation’s cortical areas and major subcortical structures.

Figure 17 shows volumetric CBF and ATT maps for a single subject of the HCD cohort. Though these appear to be noisy compared to typical ASL-derived maps, it should be noted they are at unusually high resolution and have not had any form of smoothing applied. Partial volume corrected variants of these maps are given in Supplementary Figure 5 and Supplementary Figure 6.

**Figure 17:**
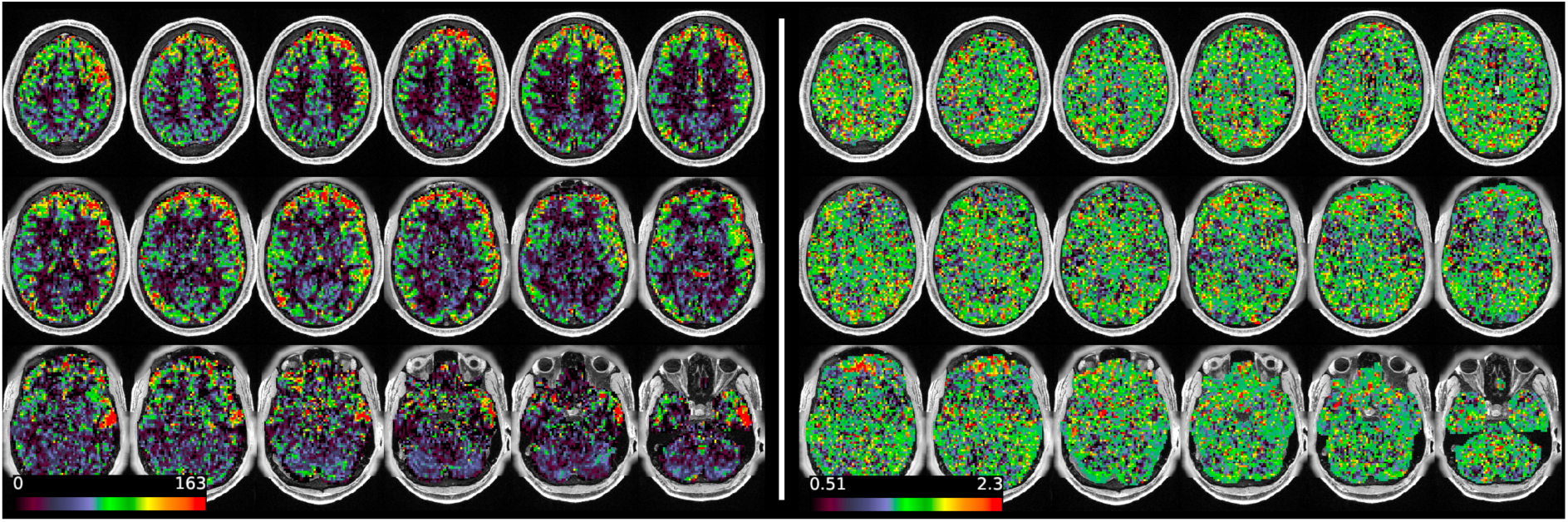
Volumetric CBF (left) and ATT (right, both non-PVEc) maps for subject HCD0378150 of the HCD cohort.

## 4. Discussion

### 4.1. Haemodynamic measures: CBF and ATT

The HCP Lifespan ASL dataset and accompanying pipeline have yielded a dataset that is unusually large and sophisticated compared to existing ASL studies. The use of a high spatial resolution multiple delay acquisition has enabled CBF and ATT maps to be produced in both volumetric and surface representations. The group average surface-based ASL measures, which were generated using areal-feature-based surface registration with minimal smoothing, represent the most high-resolution maps of brain perfusion currently available, and will enable the wider exploration of surface-based analysis techniques for ASL.

CBF on the cortical surface was often observed to follow cortical areal boundaries (see Figure 7 and Figure 9), which suggests that cerebral perfusion may be further regulated within vascular territories. Given that prior results from HCP fMRI data have established functional and/or connective relationships between cortical areas, it would be interesting to investigate whether perfusion is a further component of these relationships. Furthermore, given perfusion is a component of the BOLD effect, greater understanding of this parameter in the cortex will be of utility to fMRI studies. Comparing between cohorts, the HCD cohort had higher CBF than HCA (a difference of 24.7 ml/100g/min), though the difference was slightly reduced by PVEc (23.4 ml/100g/min), which is consistent with age-related atrophy leading to increased PVE presenting a confounding factor for perfusion measurement with ASL on elderly cohorts.

The ATT maps, both surface and volumetric, evidenced longer transit times in posterior portions than the anterior portions, which is expected because the vascular calibre of the common and internal carotid arteries that feed the ACA and MCA is larger than that of the vertebral arteries that feed PCA through the basilar artery, resulting in faster flow at a given blood pressure. Further, the HCA cohort showed higher ATT than HCD (by 0.2s), which is consistent with previously reported findings that ATT increases with age. The group mean volumetric ATT maps showed some small differences in GM/WM contrast between the two cohorts. One explanation for this could be reduced contrast in core/watershed ATT in the HCD cohort, making GM/WM contrast more prominent, whereas in the HCA cohort core/watershed contrast dominates, possibly due to vascular disease.

Due to the high spatial resolution of the acquisition, the volumetric maps produced by the pipeline appeared noisier than ASL acquired at typical resolution (e.g. 3.4 x 3.4 x 5mm). This was particularly the case in WM (shown for example in Figure 17, Supplementary Figure 5 and Supplementary Figure 6). Not only does WM have intrinsically lower perfusion (and thus SNR) than GM, which poses a fundamental challenge for ASL, but the acquisition itself was optimised for the cortex (high spatial resolution was used to facilitate accurate registration to the cortical surface, and the sensitivity profile for the multi-channel coil used preferentially increased SNR in the cortex). Under the Bayesian framework used by the FSL BASIL toolbox for perfusion quantification, parameter values will revert to their prior mean values when the data is uninformative. For WM ATT voxels this is a likely explanation for why values tend to be near the prior mean of 1.6s, although this would be expected to only have a small effect on the estimated perfusion value. Ultimately the pipeline presented here has not been optimised for WM voxels and more precise results might be achieved by averaging over regions in WM.

Nevertheless, recent work investigating high resolution ASL has found it to be practical and feasible for perfusion measurement in the cortex and subcortex, despite the appearance of noise, because the reduction of inherent partial volume effects improves the localisation of perfusion without impacting sensitivity (Kashyap et al., 2024). A conventional mitigation for noise would be to perform spatial smoothing (similar to reducing spatial resolution), at the loss of spatial precision. The alternative strategy adopted here is to average perfusion measurements within neuroanatomically well-defined subdivisions to make efficient use of available SNR. This should increase statistical sensitivity for effects, while reducing dimensionality (thereby reducing the penalty for correction of multiple comparisons) without the need for indiscriminate spatial smoothing. These imaging-derived phenotypes should facilitate novel statistical analyses such as relating CBF and ATT to behaviour, cognition, genetics, or clinical conditions.

In both cohorts, PVEc was observed to lead to statistically significant increases in CBF estimates. This is despite the relatively high spatial resolution of the ASL data (2.5mm isotropic), which reduces but does not eliminate the presence of PVE in the data. Indeed, Figure 13 showed that PVEc led to greater increases in cortical than subcortical CBF, which is consistent with there being more PVE present in cortical voxels. Recent work has noted the fundamental trade-off inherent to PVEc, while acknowledging that both PVEc and non-PVEc data have value depending on the research question (Chappell et al., 2021). Without PVEc, differences in GM PVE between individuals due to differences in anatomy are encoded into the perfusion values, thus becoming a source of between-individual variability. Though PVEc mitigates this, it also increases the complexity of the model by introducing tissue-specific parameters which could result in over-fitting, introducing a different source of between-individual variability.

Both the data acquisition and processing pipeline used in this work were optimised for producing surface maps of perfusion.

### 4.2. Correction of acquisition artefacts

The novel nature of the multi-band acquisition and the high demands for precision in spatial localization in the HCP-Style approach to data acquisition and analysis (Glasser, Smith, et al., 2016) necessitated careful pre-processing of the ASL and calibration data to bring it into T1w alignment space at the original voxel resolution with properly normalized image intensities. The pipeline that has been implemented represents a best-attempt at preprocessing given the unique nature of the acquisition, though it has not been compared against existing ASL pipelines. Recognising that users may wish to implement their own alternative kinetic modelling to the variational Bayesian method used in this work, the HCP-ASL processing pipeline also provides fully-corrected ASL and calibration volumetric data, which means that a bespoke perfusion estimation may be performed without needing to re-create the prior correction steps.

Addressing the SMS banding artefact was a particular pre-processing challenge. The banding remaining after correction for saturation recovery is suspected to be caused by magnetisation transfer effects, namely that the acquisition of one slice has spill-over effects on the adjacent slice acquired next. In the derivation of the empirical correction used in this work, it was found that the coefficients differed slightly between samples of the HCA and HCD cohorts (illustrated in the Supplementary Material). This could be due to age-related differences in grey matter and white matter volumes. Ultimately it was decided to use a single set of correction coefficients derived over an equal-sized sample of both cohorts for simplicity, but in the future a personalized correction technique to remove the need for a representative population might be feasible and would allow the pipeline to translate to unseen datasets with ease. A personalised approach could also be beneficial for application to subjects where tissue T1 may be modified by pathology, as this is mechanism at least partially explains the banding artefact.

Across both cohorts, lower CBF was observed in the most inferior cortical regions such as the inferior temporal cortex and orbito-frontal cortex. It is likely that this was caused by uncorrectable susceptibility-induced signal loss. Though distortion correction can relocate signal to the correct position within the brain, it cannot correct the underlying loss of signal, which could only be solved by the use of a spin-echo acquisition.

### 4.3. Relationship of ATT to fMRI temporal components

In previous work, group temporal independent components were generated from the HCA and HCD blood oxygenation-level dependent functional MRI (BOLD fMRI) data using independent component analysis (Glasser et al., 2018, 2019). This revealed a global respiratory signal, driven by fluctuations in pCO2 that lead to global fluctuations in blood flow, e.g., (Power et al., 2020), which acts as an endogenous contrast mechanism leading to global blood flow fluctuations that result in global BOLD signal contrast fluctuations (i.e., global brightening and darkening of the entire grey matter). Some of the temporal independent components have been termed “vascular delay” components because they are assumed to represent the portion of global blood flow fluctuations resulting from variation in blood pCO2 that is either ahead of the primary venous phase (core regions of vascular territories with fastest perfusion) or trails the primary venous phase (watershed regions between vascular territories). The vascular delay components are expected to correlate with ATT derived from ASL, indeed, ATT-like maps have been directly produced from fMRI using vascular lag models (Tong et al., 2017, 2019).

As an exploratory example of what is feasible having surface-based perfusion measures from ASL in the HCP data, this hypothesis was tested using the derived ATT maps by calculating spatial correlation coefficients with the group-average vascular decay temporal ICA components in CIFTI greyordinates space, shown in Figure 19 and Figure 18 for the two components with the highest correlations. Whilst both cohorts showed similar spatial correlation coefficients in the first component (0.46 and 0.40 for the HCA and HCD respectively), they differed in the second component (0.51 and 0.69 respectively), though it should be noted that the second tICA component itself differed more substantially between the two cohorts (for example, the HCD cohort showed more negative values than HCA in core vascular territories).

**Figure 18:**
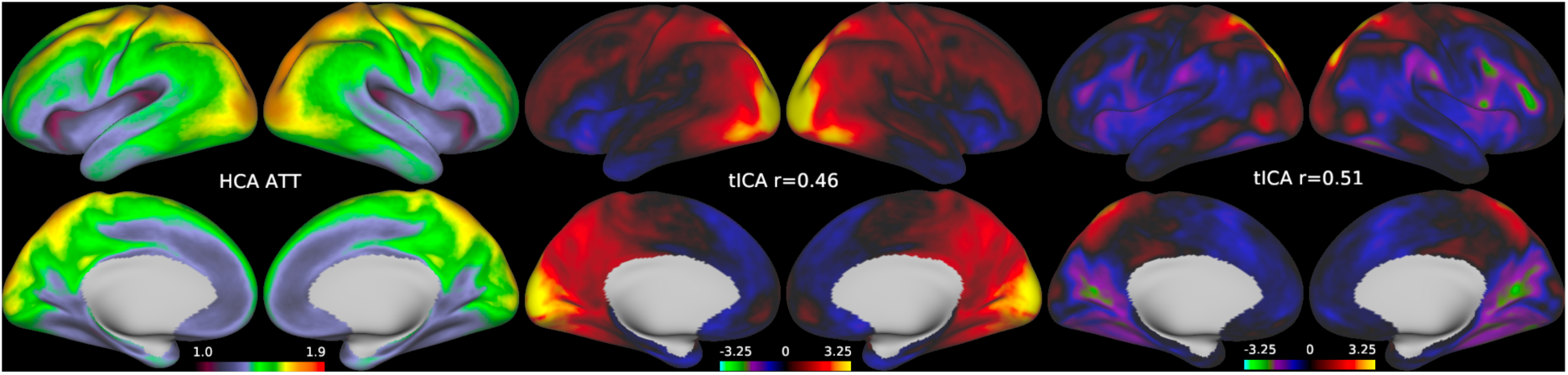
Left: Group mean PVEc ATT (sec) for the HCA cohort. Middle and right: the two most correlated tICA components (in percent of BOLD signal) for the HCA cohort, with the spatial correlation with the ATT map indicated. Only surface greyordinates are shown, though the correlation calculation included subcortical grayordinates.

**Figure 19:**
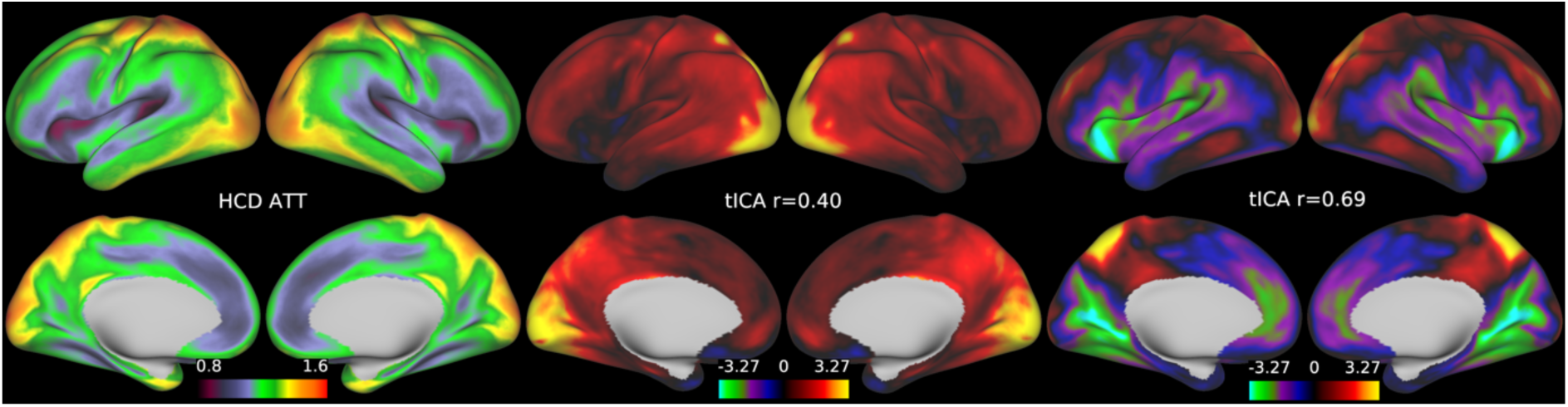
Group mean PVEc ATT (sec) for the HCD cohort. Middle and right: the two most correlated tICA components (in percent of BOLD signal) for the HCD cohort, with the spatial correlation with the ATT map indicated. Only surface greyordinates are shown, though the correlation calculation included subcortical grayordinates.

### 4.4. Further work

One area in which the pipeline could be further improved is denoising using independent component analysis (Carone et al., 2019). Preliminary work with this approach was unable to reliably identify noise components so it was not adopted for the final pipeline. This could be due to the relatively limited number of timepoints in ASL data relative to fMRI, which reduces the performance of ICA (i.e., ICA’s performance is markedly improved in high spatial and temporal resolution fMRI relative to legacy low spatial and temporal resolution fMRI). A further area of future development would be to adopt a fully surface-based approach to perfusion estimation. Since the BASIL toolbox operates in volume space, currently the pipeline produces all perfusion estimates in voxel space and then projects these onto the cortical surface. A more direct, and potentially more advantageous, approach would directly estimate perfusion on the cortical surface after projecting the ASL timeseries from volumetric data. This would allow spatial regularisation (which is beneficial given the inherently low SNR of ASL data) to be applied in a manner that respects the existing HCP principle of constrained smoothing within neuroanatomically well-defined tissues or brain areas. Indeed, one could in theory directly model the ASL parameters within brain areas, a technique that has previously been shown to be beneficial for task-based fMRI GLM modelling (Glasser, Coalson, et al., 2016).

## 5. Conclusion

The HCP Lifespan ASL dataset is a large and high-quality resource that will enable perfusion to be studied in unprecedented detail during the development and ageing phases of life. The HCP-ASL pipeline has been developed as a best-attempt to implement an HCP-Style data analysis whilst accounting for the specific nature of the acquisition with state-of-the-art correction techniques. Perfusion estimation has been performed on the pre-processed data using an established variational Bayesian method, though investigators can run their own perfusion estimation on the pre-processed data if they wish. Given ongoing debate on the utility of partial volume correction, particularly where ageing and pathology are concerned, both corrected and non-corrected perfusion estimates have been produced to provide investigators maximum flexibility. A preliminary group analysis revealed expected differences in haemodynamics between the Development and Ageing cohorts, specifically, reduced CBF and elongated ATT with increasing age.

## Acknowledgements

JT and MAC have received support from the Engineering and Physical Sciences Research Council UK (EP/P012361/1, EP/P012361/2). FKM and MC were supported by the Beacon of Excellence in Precision Imaging, University of Nottingham. YS is supported by the Royal Academy of Engineering under the Research Fellowship scheme (RF/201920/19/236). The Wellcome Centre for Integrative Neuroimaging is supported by core funding from the Wellcome Trust (203139/Z/16/Z). MPH is supported by grants: U01MH109589 and U01AG052564. MFG receives support from grants: R24MH108315, R24MH122820 and U19AG073585. TFK was funded by the Bellhouse scholarship at Magdalen College, Oxford.

## 6. Ethics

Ethical approval was secured during data collection prior to this work (https://doi.org/10.1016/j.neuroimage.2018.09.060); further approval was not required for this work.

## 7. Data and code availability

The pipeline code may be found at https://github.com/physimals/hcp-asl/releases/tag/v0.1.2 and the pre-processed data will be available at the NIH data archive (NDA): https://nda.nih.gov

## 8. Author contributions

TFK, FAKM and JT developed the pipeline code and prepared the manuscript. MSC contributed to pipeline code. DC, XL and YS contributed to methodology and reviewed the manuscript. TSC contributed to pipeline code and reviewed the manuscript. MPH and MFG contributed to methodology, performed data analysis, and reviewed the manuscript. MFG also secured funding. MAC supervised pipeline development, contributed to methodology, secured funding and reviewed the manuscript.

## 9. Author contributions

The authors declare no competing interests.

## Supplementary material

### Data acquisition sites

**Supplementary Table 1:** number of subjects acquired at each site.

|  | Harvard University | Massachusetts General Hospital | University of California, Los Angeles | University of Minnesota | Washington University, St Louis |
| --- | --- | --- | --- | --- | --- |
| HCD | 352 | 0 | 344 | 318 | 292 |
| HCA | 0 | 293 | 298 | 307 | 301 |

The distribution of data acquisition across sites is given in the below table.

### Saturation recovery T1 estimation

Supplementary Figure 1 shows an example T1 map estimated by FSL FABBER used to perform saturation recovery correction. The contrast in GM / WM T1 values means that these tissues will have been differentially affected by the saturation recovery effect, and thus require different amounts of correction which is achieved by using a voxel-wise map of T1 estimates.

**Supplementary Figure 1:**
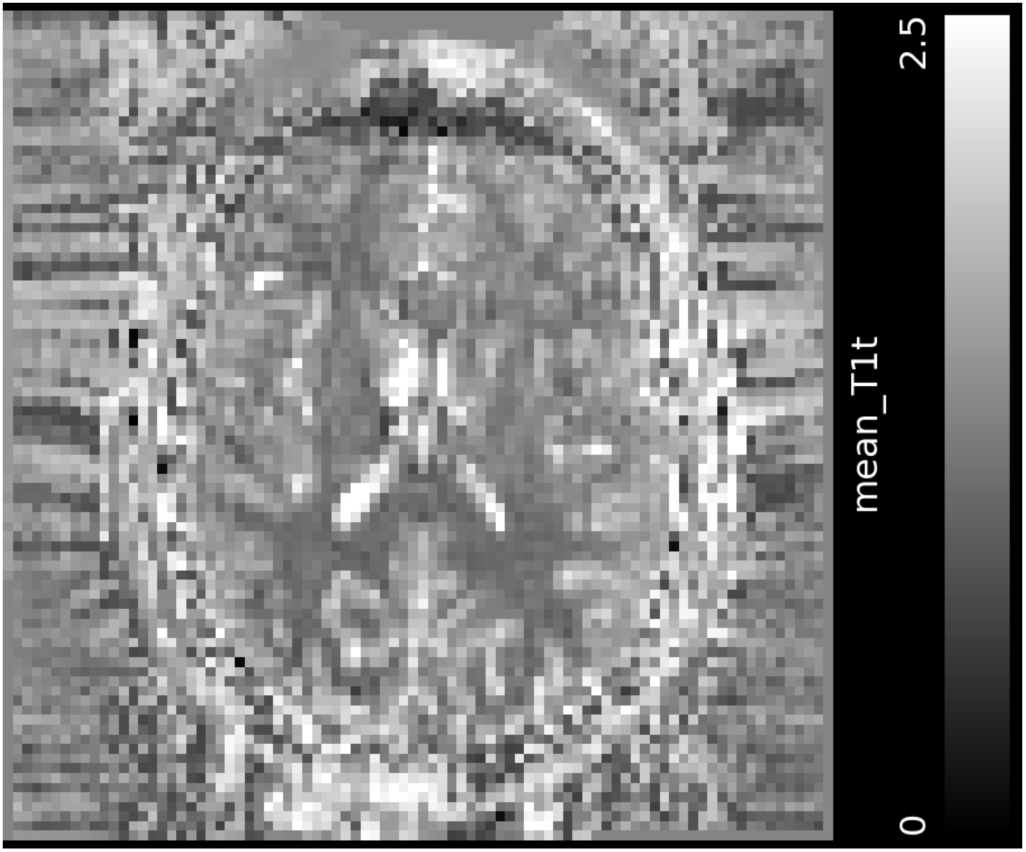
Example T1 map (axial slice) generated by FABBER for a single subject, used for saturation recovery banding correction (units in seconds).

### Derivation of the empirical banding correction

Saturation recovery correction alone was found to be insufficient to fully remove the banding artefact from the ASL and calibration data. The remaining banding is thought to arise from magnetisation transfer effects, for which separate empirical correction factors were derived in the following manner using 40 randomly selected participants from each of the Aging and Development cohorts.

First the M0 images were corrected for saturation recovery and bias field effects as described in the Methods. Grey matter (GM) and white matter (WM) FreeSurfer segmentation masks were taken from the structural pre-processed outputs of each participant’s T1w anatomical image and linearly registered with the corresponding M0 images using the registrations derived earlier in the pipeline. In the space of the M0 images, the GM and WM masks were unified to produce a brain tissue mask, which was used to calculate the mean signal intensity within each slice of the M0 image. The slice-wise mean signal intensities were averaged across the subset of each cohort to generate a slice-wise population-averaged mean M0 signal intensity profile. The profiles for each cohort (given in Supplementary Figure 2) showed an approximately linear decrease in M0 signal intensity over the 10 slices in each of the central four slice stacks. The magnitude of this effect relative to the signal intensity of the first slice in each band was estimated by fitting a single linear function to the mean of the central four bands. Correction was then performed by multiplying the image data in each band by the inverse of the linear function. The signal intensities in the two outermost (inferior and superior) bands were not used to calculate the banding correction due to the low number of tissue voxels in these bands leading to high variability, though the fit derived from the central four bands was extrapolated to cover all six bands when performing the correction. Finally, the mean of the coefficients derived in each cohort was taken to give a single overall set of corrective coefficients. Supplementary Figure 3 gives an schematic of this process.

**Supplementary Figure 2:**
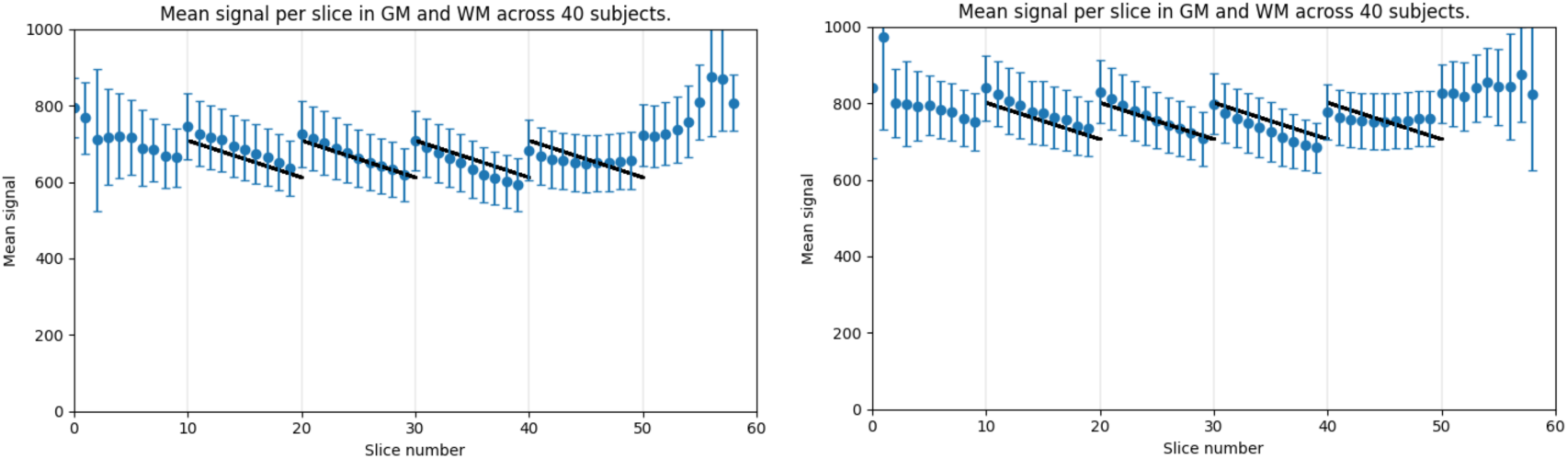
Mean calibration image signal in WM and GM by slice number for 40 HCA and HCD subjects.

**Supplementary Figure 3:**
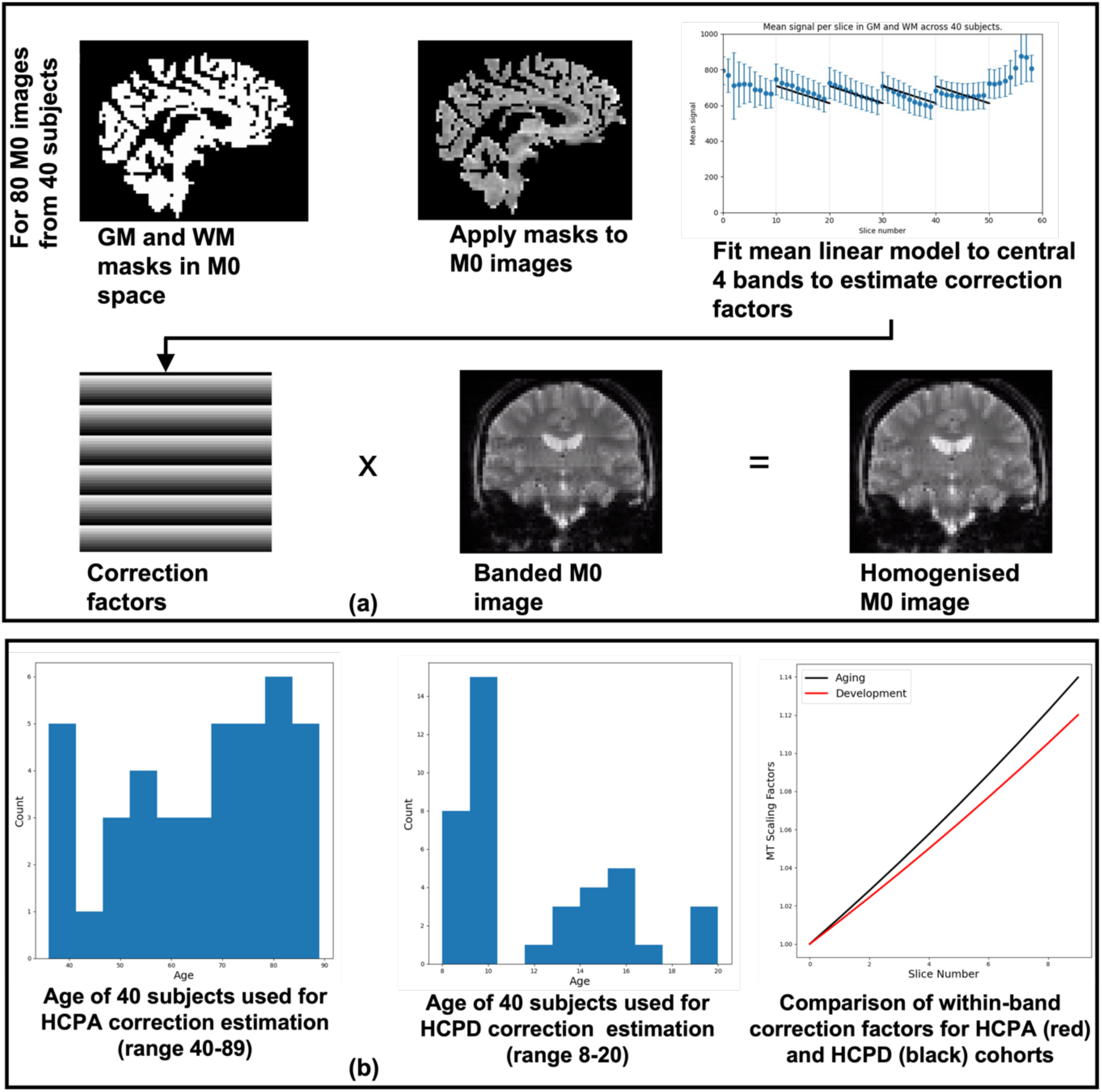
Process used to produce population derived scaling factors to correct for slice-wise intensity variations (a), thought to arise from an MT effect resulting from the use of a SMS acquisition to achieve high resolution for the HCP Lifespan ASL data. (b) shows the age distributions of the 40 subjects used to derive correcting factors for the Ageing (left) and Development (centre) cohorts. A comparison of the two sets of scaling factors is shown on the right of (b).

### Comparison of standard and boundary-based registration (BBR)

Supplementary Figure 4 shows registration between the calibration image and cortical surfaces (in T1w alignment). The initial registration is performed directly between the calibration and T1w image; for the final registration, BBR has been used between the ASL space CBF map and T1w image to refine the registration, which is then combined with the earlier motion correction matrices to obtain a new calibration/T1w registration. The difference between the two registrations is small, but slightly better for the final (refined) registration using the CBF map.

**Supplementary Figure 4:**
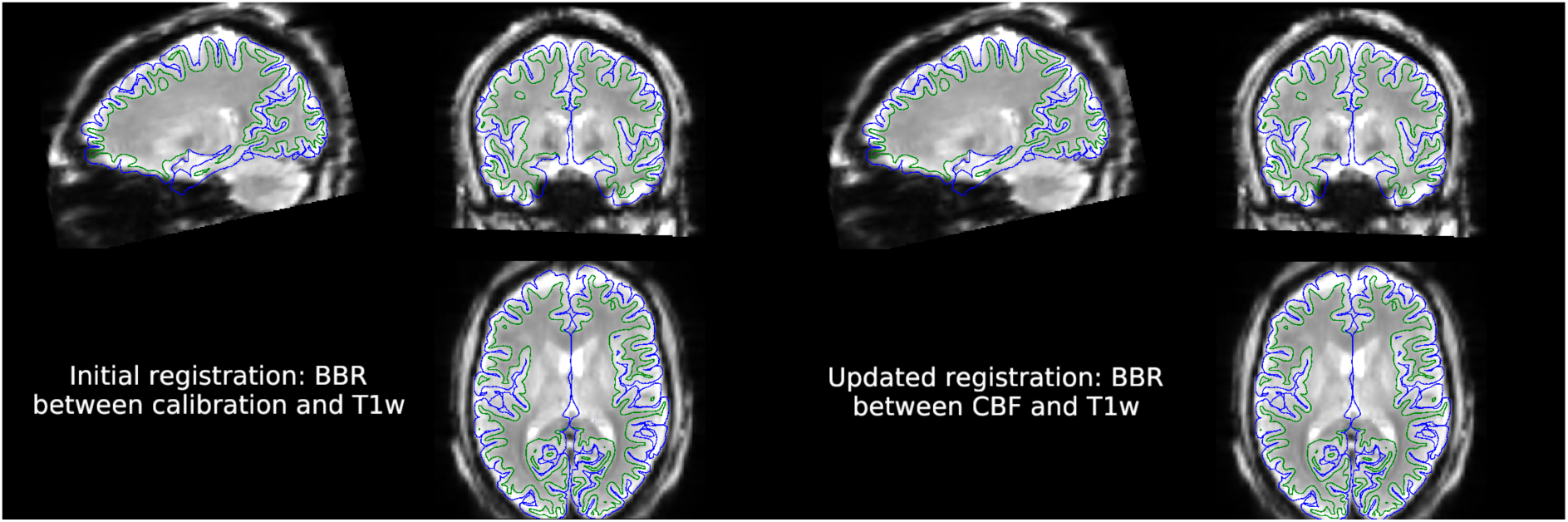
Comparison of registration initially (left) and after final (right) registrations. The calibration image is shown in alignment with the cortical surfaces in T1w space. The initial registration is obtained using BBR between the calibration and T1w; at the second step, BBR between the CBF map and T1w has been combined with motion correction between the ASL and calibration image to yield an updated calibration/T1w registration. The difference in the translation between the two registrations is less than 1mm in each dimension.

**Supplementary Figure 5:**
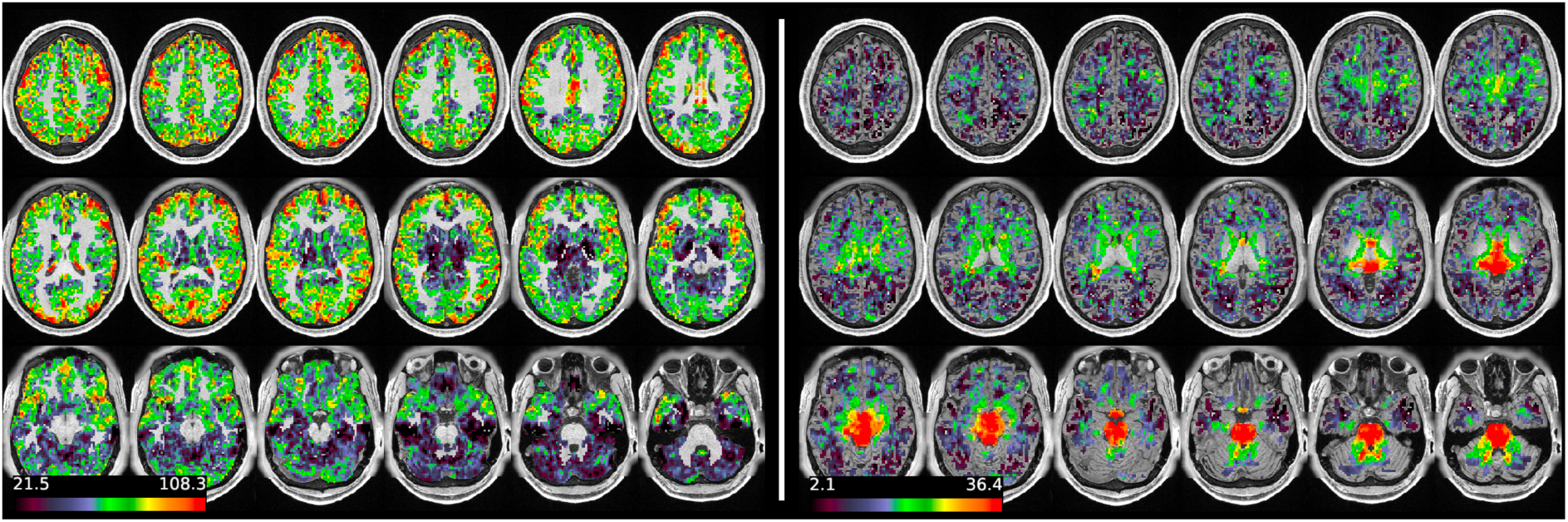
Partial volume corrected CBF maps in GM (left) and WM (right), masked for voxels with >10% of each tissue, for subject HCD0378150 (chosen at random, and also presented in the main text).

**Supplementary Figure 6:**
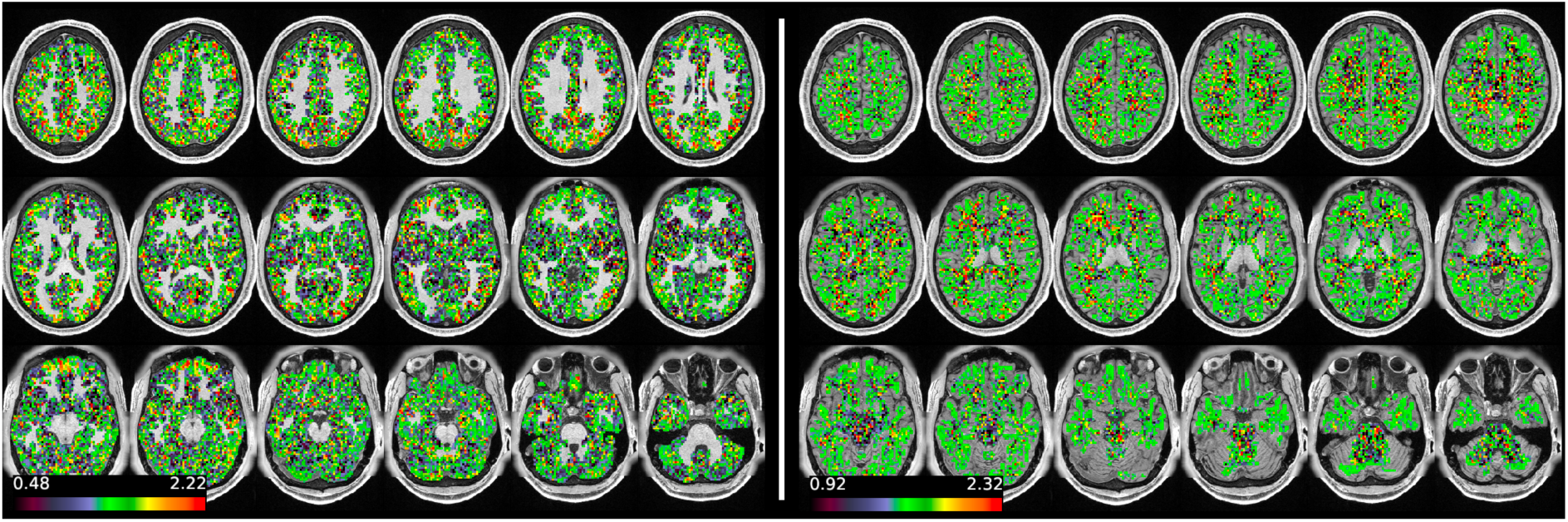
Partial volume corrected ATT maps in GM (left) and WM (right), masked for voxels with >10% of each tissue, for subject HCD0378150 (chosen at random, and also presented in the main text).

